# Multivariate climate change, the climate niche, and the Holocene history of eastern hemlock (*Tsuga canadensis*)

**DOI:** 10.1101/548420

**Authors:** Bryan Shuman, W. Wyatt Oswald, David R. Foster

## Abstract

Forests in the eastern North America have changed progressively over the 11,700 years of the Holocene Epoch. To understand the dynamics involved, we focus on eastern hemlock (*Tsuga canadensis*), which shifted its distribution through time and, notably, exhibited a rapid range-wide decline at 5280±180 YBP. We consider how climate could have shaped this history by comparing fossil pollen records from eight New England sites with quantitative temperature and effective precipitation reconstructions and evaluating the realization of *Tsuga*’s climate niche through time. The comparisons indicate that multivariate climate change significantly influenced *Tsuga* abundance, including its abrupt decline and recovery. The comparisons show that the realized climate niche of *Tsuga* expressed today includes two important features that persisted through time. First, *Tsuga* pollen percentages reach their maxima (>30%) where July temperatures equal 18-20°C, but do so at two modes where annual precipitation equals either ∼1100 or ∼775 mm. The bimodality reflects *Tsuga*’s two geographic modes in the Great Lakes and Appalachian regions today, and explains past dynamics, such as short-lived peaks in *Tsuga* abundance associated with effective precipitation of ∼775 mm at ca. 10,000 years before CE 1950 (YBP). Second, the two peaks in *Tsuga* abundance follow negative correlations between temperature and precipitation such that the two modes shift toward high precipitation if temperatures are low (e.g., ∼1400 and ∼1000 mm at <18°C). Consequently, rapid cooling at 5200±100 YBP facilitated widespread *Tsuga* declines because cooling did not coincide with increased precipitation. Abundance declined as local climates departed from optimal temperature and precipitation combinations. Recovery only followed as effective precipitation increased by >150 mm over the past 4000 years. A regionally calibrated model of the relationship of *Tsuga* pollen percentages to temperature and precipitation explains 70-75% of the variance in the percentages at eight study sites. Iteratively excluding each site from the model shows that accurately representing the major features of the climate niche enables the model to predict the mid-Holocene decline and other past changes at the excluded site (site-level RMSE = 2.1-5.6%). Similar multivariate climate dynamics closely modulated the species’ abundance throughout the Holocene with no evidence of additional large-scale disturbances.

## 1. Introduction

Climate change, broad-scale disturbances, and other factors can alter the distributions of tree species in a variety of ways, but anticipating future change is difficult (Allen et al., 2010; Lenoir and Svenning, 2014). Useful precedents exist, however, in the Holocene history of eastern North American tree species such as *Tsuga canadensis* (L.) Carrière (eastern hemlock). Today populations of this shade-tolerant, long-lived canopy tree grow in dense stands on moist, acidic or rocky soils from the Great Lakes to the Appalachian Mountains (Foster, 2014)(Fig. 1A). They can occupy multiple contrasting habitats on landscape scales (Kessell, 1979), and have different local adaptations to warm, dry versus cool, wet climates (Eickmeier et al., 1975). Such a negative trade-off between temperature and moisture may shape its distribution. Populations have expanded into new dry sites that were warm, as well as wet sites that were cool (Calcote, 2003). Populations appear limited by temperature in some areas (e.g., mountainsides in New Hampshire; Davis et al., 1980) and moisture in others (e.g., Michigan; Brubaker, 1975). Consequently, *Tsuga*’s history represents a model system for studying biogeographic responses to temperature and moisture changes, especially because the taxon’s range, abundance, and association with other taxa changed continuously as climate changed since the last ice age (Davis, 1981; Davis et al., 1986; Graham and Grimm, 1990).

**Figure 1.**
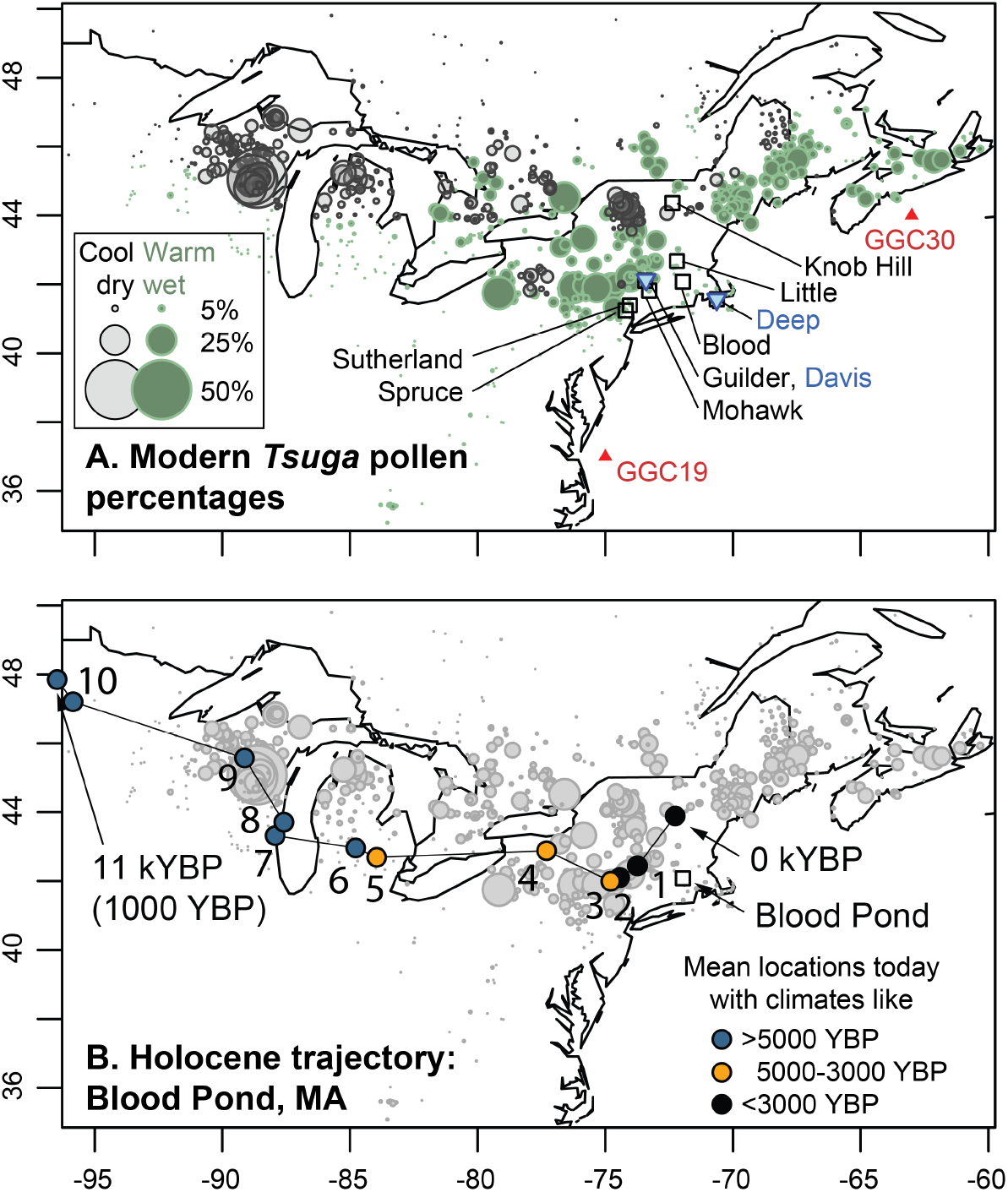
A) Map of the percentages of *Tsuga* pollen in modern sediment samples (gray and green circles) from Whitmore et al. (2005) and the locations of records used in this study (squares, pollen; blue triangles, lake levels; red triangles, sea-surface temperatures). Gray circles represent samples from cooler and drier sites than the mean of all samples with >1% *Tsuga* pollen; green circles indicate warmer and wetter sites than the mean. B) Filled circles indicate where plant populations would have to grow today to experience climates equivalent to those reconstructed for the past at Blood Pond, Massachusetts (square), labeled by millennium before present (kYBP). The west-to-east trajectory of analogous climates passes into and out of regions with abundant *Tsuga* pollen based on the mean latitude and longitude of all modern pollen samples with July temperatures and annual precipitation within 0.5°C and 50 mm respectively of past conditions inferred from alkenone paleothermometry (Fig. 2A; Sachs, 2007) and lake-level data (Fig. 2B; Newby et al., 2014). Symbol colors represent the time period represented. Gray circles represent the modern percentages of *Tsuga* pollen as in A.

Fossil pollen records spanning >12,000 years indicates that the distribution of *Tsuga* shifted northward from the central Appalachians into the northeastern U.S. (hereafter “New England”), and then westward into the northern Great Lakes region (Davis et al., 1986; Webb III, 1988; Parshall, 2002; Williams et al., 2004). The long-term geographic shifts have been widely attributed to climate changes (Fig. 1B, 2A-B)(e.g., Prentice et al., 1991; Shuman et al., 2002), but *Tsuga*’s abundance also declined precipitously across its range at ca. 5700-5000 calendar years before CE 1950 (“YBP”) and remained low for many centuries to millennia before recovering in some regions (Davis, 1981; Allison et al., 1986; Bennett and Fuller, 2002). The decline played out within decades or less at some sites (Allison et al., 1986) and only rarely involved other taxa (Foster et al., 2006). Its association with climate change has not been clear.

In one eloquently articulated hypothesis, the rapid decline resulted from biotic interactions, such as a disease or insect outbreak, in the absence of synchronous rapid climatic change (Davis, 1981; Booth et al., 2012). The rapidity of the decline, the strong single-species dynamic, and evidence of forest insect outbreaks in a few locations at the time have provided enduring support for this hypothesis (Allison et al., 1986; Anderson et al., 1986; Bhiry and Filion, 1996). However, insect outbreaks did not affect all sites and other tree species with distinctly different ecologies also declined in some areas (Foster et al., 2006; Wang et al., 2015; Oswald et al., 2017). The primary alternative hypothesis links the decline to climatic change (Deevey, 1943; Yu et al., 1997; Haas and McAndrews, 1999; Foster et al., 2006; Shuman et al., 2009; Zhao et al., 2010). Drought has usually been regarded as the primary driver, but some evidence also points to a role for low temperatures (Calcote, 2003; Marsicek et al., 2013). Given the alternatives, the decline could represent an epiphenomenon initiated by one factor and then sustained by another (Booth et al., 2012).

Several enigmatic elements could hold the key to diagnosing the causes (Fig. 2). For example, low *Tsuga* abundance after the decline persisted as long as drought was sustained (e.g., Booth et al., 2012; Marsicek et al., 2013), but effective precipitation (the net balance of precipitation and evapotranspiration) during the associated droughts was not as low as during earlier periods when *Tsuga* was highly abundant (Fig. 2B). Additionally, not all droughts associated with the decline began at the same time (Fig. 2B)(Newby et al., 2014), and in Michigan specifically, *Tsuga* pollen percentages declined before drought intensified (Booth et al., 2012). Other unexplained aspects of the decline include why other taxa sometimes declined locally, but never across their entire ranges (Davis, 1981; Foster et al., 2006); why *Tsuga* did not shift towards a region with a more suitable climate (Shuman et al., 2009); and what caused an earlier peak and decline at ca. 9800 YBP in some areas (Zhao et al., 2010)(Fig. 2C).

**Figure 2.**
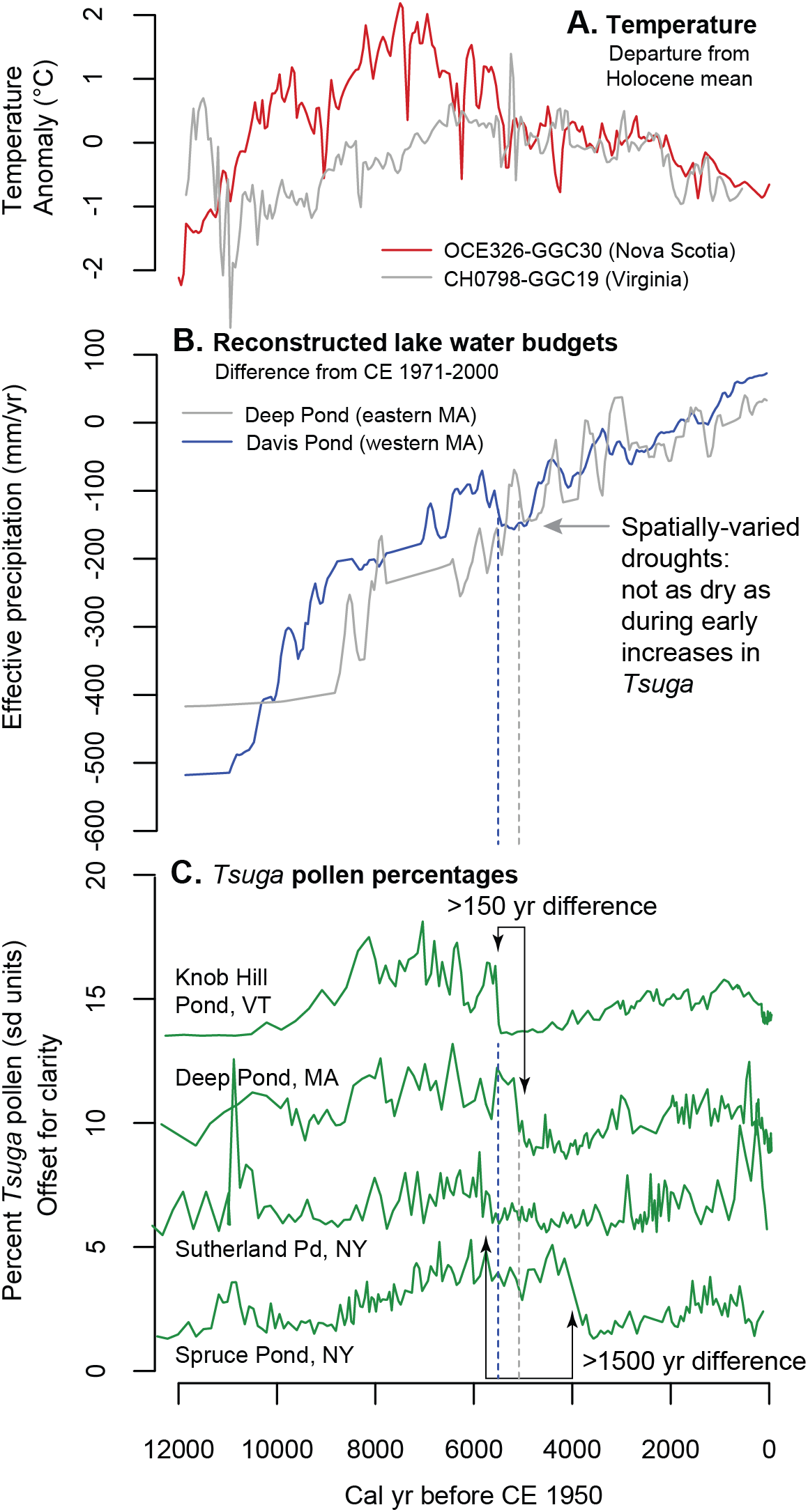
A) Holocene temperature changes reconstructed from alkenones from ocean cores GGC30 off Nova Scotia (red line) and GGC19 off Virginia (gray line)(Sachs, 2007). B) Effective precipitation reconstructions calculated from lake-level changes at Davis (dark blue) and Deep ponds (gray)(Marsicek et al., 2013; Newby et al., 2014). C) Representative *Tsuga* pollen time series (normalized to z-scores for visualization) with arrows highlighting enigmatic features of the record, including differences in decline timing.

Finally, the *Tsuga* decline can no longer be assumed to be either a widely synchronous event or the result of widespread rapid mortality as was originally suspected (Davis, 1981; Webb, 1982). Differences in both the timing and rate of decline have not been satisfactorily resolved, and some declines could represent a lack of regeneration and replacement rather than abrupt mortality (Fig. 2C). Differences exist even in records with limited temporal uncertainty and high sample density (see Liu et al., 2012), such as Spruce and Sutherland ponds in New York’s Hudson Highlands (Maenza-Gmelch, 1997; Shuman et al., 2009)(Fig. 2C). The mean age of the decline in New England equals 5280±180 YBP (Shuman et al., 2009), but the decline at Spruce Pond dates to 3800±100 YBP (Maenza-Gmelch, 1997). These differences raise doubts about explanations such as rapidly spreading diseases or insect outbreaks.

Here, we evaluate the interactions of *Tsuga*’s multi-dimensional climate niche with the multivariate array of Holocene climate changes (Shuman and Marsicek, 2016). Environmental or Grinnelian niches often link the presence or absence of a taxon to factors such as climate (Peterson et al., 2011), but here, we focus on how abundance, represented by pollen, relates to climate. Non-linearities and multidimensionality within the niche could have produced counter-intuitive outcomes, especially when combined with the different independent trajectories of both temperature and moisture (Webb III, 1986; Crimmins et al., 2011)(Fig. 3). Changes may have played out via effects on physiological or ecological traits across different life history phases and populations (e.g., Grubb, 1977; Webb III, 1986; Davis and Shaw, 2001). From a given starting point, either temperature or moisture can independently alter abundance (e.g., bold arrows, Fig. 3A) and decreases or increases can arise where the opposite response might be expected if only one factor were considered (e.g., the hypothetical site trajectory in Fig. 3A where abundance reaches a minimum between period 1 and 2 at the model species’ moisture optimum because temperature creates marginal conditions). Anticipating change is further complicated because the niche may not be fully realized at any given place in time or space (Gaston, 2003; Broennimann et al., 2007; Nogués-Bravo, 2009; Veloz et al., 2012; Maiorano et al., 2013).

**Figure 3.**
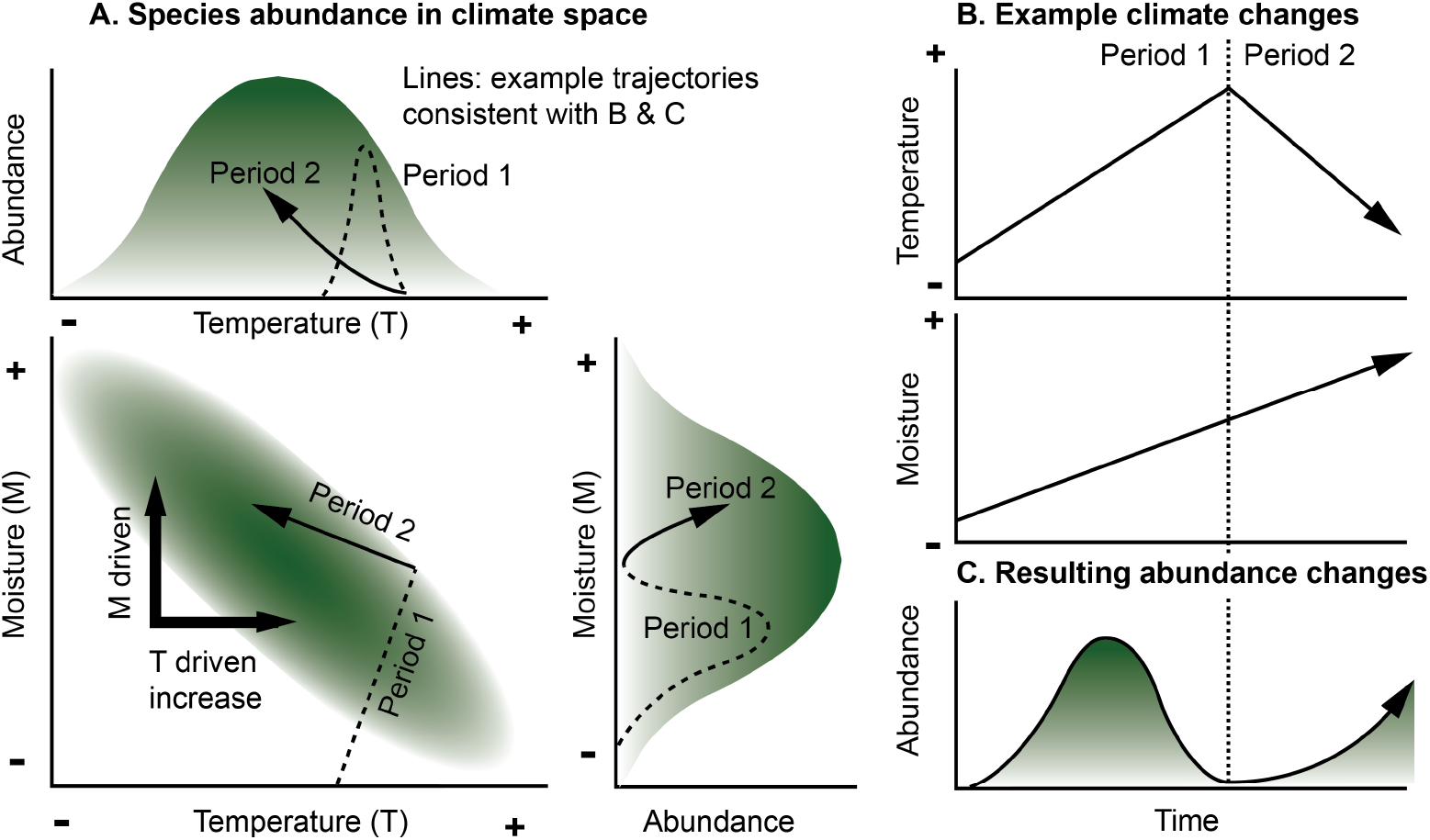
A conceptual model shows how asymmetric, multivariate climate niches can produce non-linear and unexpected responses to linear climate trends. In A, green shading represents increased *Tsuga* abundance with respect to temperature (top in A), moisture (right in A), and the interaction of the two climate variables (lower left in A). Bold arrows in A illustrate that changes in either climate variable could result in changes in *Tsuga* abundance. The thin arrow in A represents a possible trajectory through the niche based on the climate history represented for two periods in B; the dashed lines in A show where the first period would plot and the solid line shows the second period. Because of the interaction of temperature and moisture, optimal moisture conditions may coincide with minimal *Tsuga* abundance at the boundary between periods 1 and 2, and the trajectory does not project intuitively into uni-modal climate space for either temperature or moisture.

We first describe the modern realized climate niche of *Tsuga* (pollen), and then discuss regional climate changes that would have interacted with *Tsuga*’s climate preferences during the Holocene. Using quantitative reconstructions of both summer temperatures and annual effective precipitation (Sachs, 2007; Newby et al., 2014), we reconstruct the past realization of the climate niche (the total climate envelope occupied at our study sites through time; Nogués-Bravo, 2009) and use a series of statistical models to evaluate the contribution of climate to the Holocene history of *Tsuga* in New England. Overall, we address two key questions: What is the structure of the realized climate niche of *Tsuga* through space and time? How well do climate changes predict the Holocene history of *Tsuga* abundance?

## 2. Methods

### Modern pollen & climate

To describe the modern relationships among climate variables and *Tsuga* pollen abundance, we use the North American modern pollen-climate dataset compiled by Whitmore et al. (2005) and described by Williams et al. (2006). We represent abundance as a percent of terrestrial pollen, and focus on data from east of 105°W longitude. To evaluate the interaction of July temperatures and annual precipitation, we plot pollen data relative to each variable, but also consider abundance relative to the interaction among variables (Fig. 4). To do so, we calculate and then sum departures from the mean of both climate variables where *Tsuga* pollen exceeds 1% (Fig. 4B-C). To do so, we first normalize the departures by the standard deviation of each respective variable. We also sub-divide the data into two sub-groups based on whether samples represent warm-moist (e.g., Appalachian) or cool-dry (e.g., Great Lakes) areas (Fig. 1A), which are associated with negative or positive departures from the climatic mean respectively (Fig. 4B).

**Figure 4.**
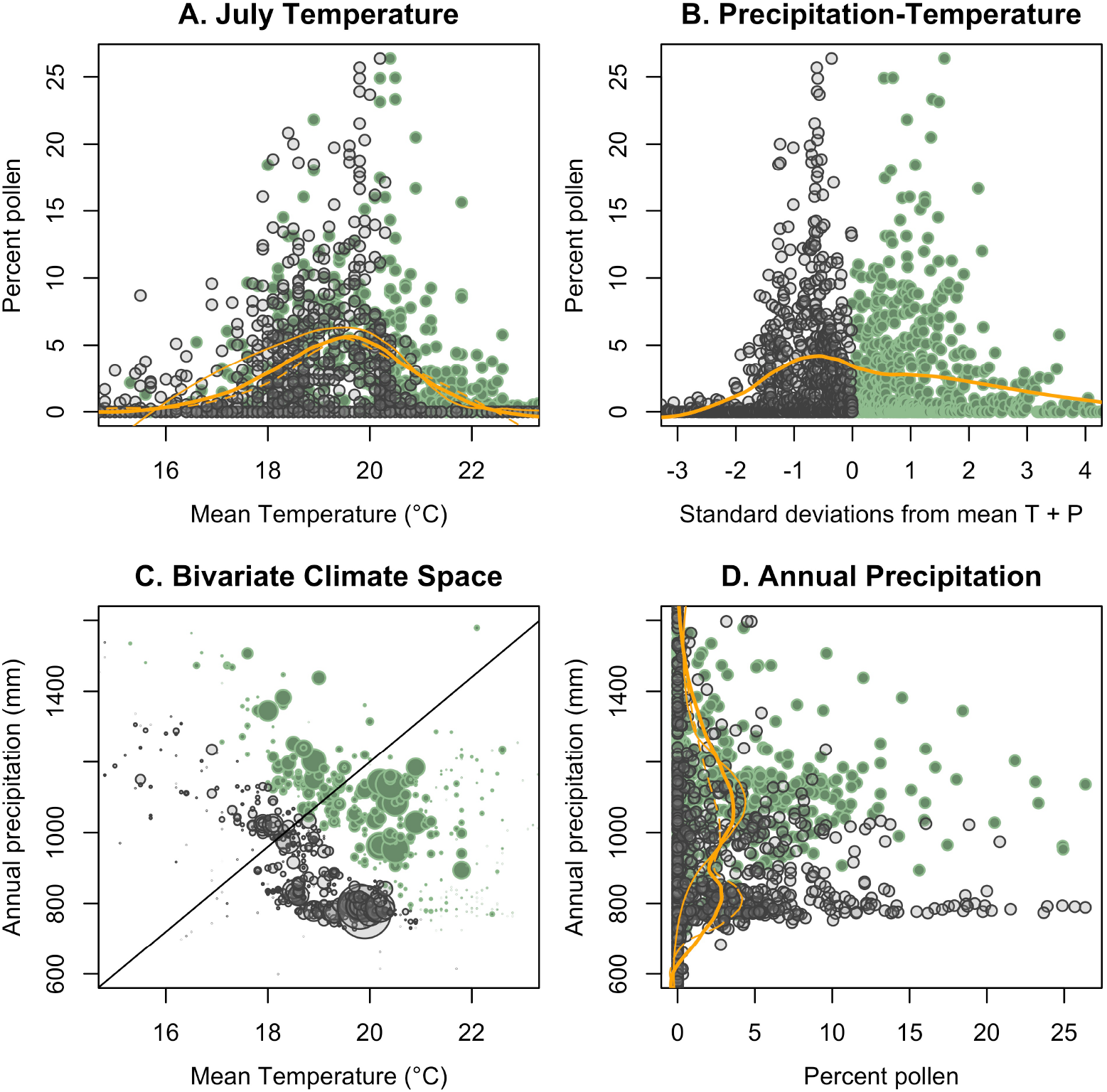
Modern *Tsuga* pollen percentages plotted relative to mean July temperature and annual precipitation. Panels A and D represent the single variable projections (sides) of the bivariate climate space shown in panel C; panel B represents a projection onto the cross-cutting diagonal line in C. Panels A and D show the percentages relative to mean July temperature (T) and mean annual precipitation (P) respectively. The x-axis in B is measured as summed standard deviations from the mean values of T and P for all samples with >1% *Tsuga* pollen. As in Figure 1, gray circles represent sites that are both cooler and drier than the mean (left side of B); green circles indicate warmer and wetter sites than the mean (right side of B). Bold orange lines show the locally-weighted means of *Tsuga* pollen percentages from all samples; dashed orange lines show the local means for cool dry samples (gray symbols) and thin solid orange lines show the local means for warm wet samples (green symbols). In C, bubble sizes represent *Tsuga* pollen percentages with largest equal to 50%.

### Fossil pollen

We evaluate past *Tsuga*-climate relationships at eight representative sites in New England (Table 1). Four sites (Knob Hill, Little, Blood, and Deep ponds) form a north-south transect of detailed pollen records spanning from cool to warm areas of New England, while a second set of four sites (Guilder, Mohawk, Sutherland, and Spruce ponds) represents pairs of high and low elevation areas in the Berkshire and Hudson Highlands of Massachusetts and New York respectively. The elevational pairs also include unusual features of the *Tsuga* pollen record, including areas where *Tsuga* populations did not recover after the decline (like in many central Appalachian sites); where *Tsuga* populations experienced two declines (at ca. 10,000 and 5000 YBP, such as at Sutherland Pond, NY, Fig. 2C); and where *Tsuga* populations declined much later than most other sites (at 3800 YBP at Spruce Pond, NY, Fig. 2C)(Gaudreau, 1986; Maenza-Gmelch, 1997). Overall, we analyzed four sites with high *Tsuga* pollen percentages (>20%) and four sites with low *Tsuga* pollen percentages (<10%). Data from most sites were generated at Harvard Forest using standard techniques (Foster et al., 2006; Oswald et al., 2007; Oswald et al., 2018; Oswald and Foster, 2012), but data for Mohawk, Spruce, and Sutherland ponds (Gaudreau, 1986; Maenza-Gmelch, 1997) were obtained from the Neotoma Paleoecology Database (Williams et al., 2018).

**Table 1:** Study sites sorted by mean July temperature. “Elev” refers to elevation; “Precip” refers to mean annual precipitation; “Dates” refers to the number of calibrated radiocarbon ages used to constrain the pollen sample ages; “Samples since 11ka” refers to the number of pollen samples in the record since 11,000 YBP (calendar years before CE 1950). “TSD” refers to *Tsuga* decline age based on the peak rate of change over 100-yr intervals rounded to the nearest 25 years; ages are presented as YBP.

| Site | PRISM 1940-1970 normals |  |  |  |  |  |  |  |  |  | Reference |
| --- | --- | --- | --- | --- | --- | --- | --- | --- | --- | --- | --- |
|  | Latitude | Longitude | Elev | July Temperature (degrees C) |  |  | Precip | Dates | Samples | TSD |  |
|  | Degrees | Degrees |  |  |  |  |  |  | Since |  |  |
|  | N | W | m | Mean | Maximum | Minimum | mm | # | 11 ka | YBP |  |
| Guilder Pond | 42.11 | 73.44 | 624 | 19.9 | 25.5 | 14.2 | 1111 | 9 | 41 | 5325 | Oswald et al., (2018) |
|  |  |  |  |  |  |  |  |  |  |  | Oswald & Foster |
| Knob Hill Pond | 44.36 | 72.37 | 370 | 19.3 | 26.3 | 12.3 | 931 | 9 | 130 | 5500 | (2012) |
| Little Pond | 42.68 | 72.19 | 302 | 20.4 | 27.5 | 13.3 | 1046 | 7 | 80 | 5300 | Oswald et al. (2007) |
| Mohawk Pond | 41.82 | 73.28 | 360 | 20.8 | 26.5 | 15.1 | 1171 | 12 | 34 | 5550 | Gaudreau (1986) |
| Blood Pond | 42.08 | 71.96 | 214 | 21.0 | 26.9 | 15.2 | 1101 | 15 | 120 | 5200 | Oswald et al. (2007) |
| Deep Pond | 41.56 | 70.64 | 23 | 21.1 | 24.9 | 17.2 | 1125 | 14 | 140 | 5150 | Foster et al. (2006) |
| Spruce Pond | 41.24 | 74.20 | 223 | 21.9 | 28.1 | 15.7 | 1162 | 9 | 141 | 3975 | Maenza-Gmelch (1997) |
| Sutherland Pond | 41.39 | 74.04 | 380 | 22.1 | 27.4 | 16.7 | 1251 | 11 | 140 | 5600 | Maenza-Gmelch (1997) |

### Climate reconstructions

We compare the fossil pollen data with quantitative temperature and effective precipitation estimates based on alkenone paleothermometry (Brassell et al., 1986; Sachs, 2007) and lake-level reconstructions (Marsicek et al., 2013; Pribyl and Shuman, 2014). These estimates derive from four locations (Fig. 1, 2): temperatures from cores collected off the coasts of Nova Scotia (GGC30) and Virginia (GGC19)(Sachs 2007) and effective annual precipitation from Davis and Deep ponds in western and eastern Massachusetts respectively (Newby et al., 2011; Marsicek et al., 2013; Newby et al., 2014). Continental temperature records are rare and difficult to obtain, but in New England, we make the assumption that regional temperature changes influenced both terrestrial and marine locations. Historically, mean annual temperatures onshore and sea-surface temperatures (SSTs) in the region correlate. Climate division data from across Massachusetts (Vose et al., 2014) for CE 1948-2016 correlate (r = 0.69) with SSTs from 42-44° N and 65-70° W (Kaplan et al., 1998); both datasets show a period of low temperatures after the warm CE 1950s and a subsequent warming trend with similar inter-annual variability. Temperature and effective precipitation time series for the Holocene also share significantly (and negatively) correlated signals at millennial to multi-century timescales despite deriving from independent data sources, which provides confidence that they represent a coordinated set of climate changes around New England (Shuman and Marsicek, 2016). Appendix I describes the specific records, their age control, and the interpolation and analytical methods.

We linearly interpolated the reconstructed climate anomalies to the locations of our pollen records based on latitude and longitude. Absolute temperature and precipitation time series for each fossil pollen record were then derived by combining the interpolated paleoclimate anomalies with the mean July temperatures and annual precipitation rates for CE 1941-1970 for each location from the 800-m resolution PRISM time-series climate dataset (Daly et al., 2008). The absolute temperature and precipitation time series, thus, account for differences related to factors such as elevation and latitude, and are important for evaluating the climate niche (Fig. 3).

### Reconstructing past climate-abundance relationships

Ancient changes in the realization of species’ climate niches have been evaluated using fossil data in combination with climate model simulations (Nogués-Bravo, 2009; Veloz et al., 2012; Maiorano et al., 2013). Here, we interpolate the paleoclimate reconstructions as the basis for our statistical model because we want to capture the specific sequence of both forced and stochastic climate variations at sub-millennial scales (Fig. 2). To match the fossil pollen percentages to the full Holocene array of mean July temperatures and annual effective precipitation for each site, we also interpolate the data in time. We use uniform 50-yr time steps to ensure that the major patterns in the original data were not aliased.

We compare current and past relationships among *Tsuga* pollen percentages, mean July temperature, and mean annual precipitation by randomly pairing modern and fossil pollen percentages from locations and times with the same climates. To do so, we subdivide all of the data into 0.5°C and 50 mm moving windows regardless of time, and compare up to 1000 random pairs per window containing at least one fossil sample and a non-zero modern median percentage (i.e., where the two realized niches overlap and are well constrained by data). In each window, if the 95% distribution of the random differences did not include zero, we infer a significant difference.

### Models of past pollen percentages

Finally, we apply generalized additive models (GAMs), which are locally weighted polynomials, to estimate how well mean July temperature and annual effective precipitation data explain the time series of pollen percentages. GAMs can account for the non-linearities in the climate-abundance relationships (Wood, 2006a). They do not fit a specific global function (e.g., linear, quadratic) to the complex relationships, but instead, use local smoothing in the reconstructed paleoclimate space (e.g. locally-weighted scatterplot smoothing or lowess; Cleveland, 1979). Our models use tensor product smooths to incorporate the interactive temperature and effective precipitation effects (Wood, 2006b). They sub-divide climate space and apply the local, non-parametric polynomials within each window in a fashion that ensures no sharp breaks in the climate-pollen relationships between windows (Wood, 2006a).

We apply the GAMs to sites individually and to the entire dataset with each site withheld iteratively for model validation. The single site (“site-specific”) models provide an estimate of how well the relative sequence of temperature and moisture changes explain the local variations in *Tsuga* pollen percentages. Because of the non-linear relationships involved, we use the GAMs in place of linear regression to measure the variance explained. We test their significance by comparing the site-specific models with a null distribution of models for each site that use only randomly generated time series with the same autoregressive characteristics as the paleoclimate reconstructions (Appendix I). If the GAMs fit the *Tsuga* pollen percentages only because of the flexibility of the GAM or the ability of similarly smoothed time series to produce spurious correlations (Granger and Newbold, 1974), the variance explained by the site-specific GAM would not exceed the range of variance that could be explained by autoregressive models alone (Telford and Birks, 2011).

The regional (“leave-one-out”) models test the relationships between *Tsuga* pollen percentages and the absolute climates rather than the relative sequence of climate changes. By iteratively excluding each site, we test the hypothesis that a broad sampling of *Tsuga*’s climate niche across time and space provides sufficient information to predict the species’ history. Because data representing multiple locations and times were required to fully sample the Holocene environmental space, the exclusion of some sites from under-sampled climate regions produced deficient models. Models excluding these sites were inadequate to test the hypothesis, but reveal characteristics of the climate niche essential for accurate prediction.

To create each regional model iteration, we used a generalized additive mixed model that combines a GAM with a linear mixed-effect model to treat the individual locations (sites) as random effects (Wood, 2006a). For each site excluded during model construction, the regional mixed-effect model included only the climate data and pollen percentages from the most recent sample to account for the random effects when predicting change earlier in the Holocene. None of the models rely on any modern calibration data, but rather determine the best fit among the paleoclimate and *Tsuga* time series either for each site individually or for the full set of calibration sites using data for all time intervals >550 BP. All of the pollen percentages were exponentially transformed by 0.25 before applying the GAM function in R (R Core Team, 2017). To further ensure that the models were not overfit, we minimized the flexibility of the smoother by restricting the maximum number of basis functions, k, for each model: 5 for site-specific models and 10 for the regional “leave-one-out” models, which incorporated more data. Doing so substantially reduced the degrees of freedom in the model and is conservative relative to the recommended values of k, which are 10n^2/9^ where n is the number of data (Kim and Gu 2004); the recommended values for our site-specific and regional models are 33 and 51 respectively.

Furthermore, we examined a multivariate climate signal akin to using sums of growing-degree days and soil moisture to predict productivity increases (Woodward, 1987; Prentice et al., 1992). Starting from the cool, dry early Holocene when *Tsuga* abundance first rose, increases in either temperature or moisture could have increased abundance (e.g., perpendicular bold arrows, Fig. 3A). To represent this heuristic climate signal, we summed the positive deviations of temperature and effective precipitation from a baseline determined by the Holocene mean at each site. The paleoclimate data were normalized to z-scores (departures from the Holocene mean in standard deviation units) and then the positive values were combined into a single time series. As long as at least one variable has a positive value, *Tsuga* abundance was predicted to be above zero (such as would be the case for either perpendicular bold arrow in Fig. 3A). We do so to further ensure that the GAM fits represent meaningful signals in the climate data rather than spurious non-linear fits. The R code and data used are provided as a Supplement to this paper.

## 3. The realized climate niche of *Tsuga canadensis* today

*Tsuga* pollen represents nearly 50% of the terrestrial pollen sum in areas with mean July temperatures of about 20°C and annual precipitation above 775 mm (Fig. 4). With respect to July temperatures, *Tsuga* pollen percentages have a unimodal and nearly symmetric distribution about a mean of 19.2°C with a standard deviation of 1.2°C (Fig. 4A). Distributions with respect to temperature are similar in both wet and dry areas (orange lines, Fig. 4A). The relationship with annual precipitation has two modes: one narrow mode near the sharp lower limit of *Tsuga* pollen percentages >1% at 775 mm and a second broad mode centered around a mean of 1106 mm (Fig. 4D). These modes correspond to peak abundances in the Great Lake region (gray symbols in Figures 1A and 4D) and Appalachian Mountains (green symbols in Figures 1A and 4D) respectively. Between the two modes at 900 mm, the percentages only achieve a maximum of ∼10% as shown by local-weighted means of the percentages with respect to precipitation (orange lines, Fig. 4D).

The two modes become more pronounced when viewed along the axis of temperature-precipitation interaction (Fig. 4B). Percentages rise to two distinct peaks of >25%: one about 1 standard deviation below the mean (cool, dry areas; negative departures from the means) and one about 1 standard deviation above the mean (warm, moist areas; positive departures from the means). When we split the data along this axis into negative and positive groups, and plot the data in bivariate temperature versus precipitation space (Fig. 4C), we observe that *Tsuga* pollen percentages increase along two parallel ridges of abundance with one representing drier and cooler sites than the other. The two groups also tend to split out geographically: the negative group predominantly corresponds to the mode of abundance in the Great Lakes region and the positive group to the Appalachian-New England mode (Fig. 1A).

In both groups, maximum abundance tracks a negative correlation between mean July temperatures and annual precipitation (Fig. 4C). Linear regression reveals that the negative group follows a line with a slope of −84.6 mm/°C and an intercept of 2480 mm (R^2^=0.66; n=1434); the positive group has a slope of −62.1 mm/°C and an intercept of 2340 mm (R^2^=0.24; n=923). Consequently, about 140 mm of annual precipitation separates the two groups, but they come close to merging together in wet and cool areas. Data gaps, which make the groups visually distinct in Fig. 4C, represent the area of the Great Lakes themselves, but low *Tsuga* pollen percentages in the trough between the groups represents the absence of extensive *Tsuga* populations in western Ohio, Indiana, southern Michigan, and other areas between the arms of the species distributions in Appalachia and the northern Great Lakes region (Fig. 1). Low abundance in this region is not a function of historic land clearance, but has persisted since before European settlement (Shane, 1991; Clark et al., 1996; Wang et al., 2015; Paciorek et al., 2016; Goring et al., 2016).

## 4. Climate, *Tsuga* abundance, and the climate niche through time

### 4.1 Climate trajectories

*Tsuga* populations in New England have experienced multiple changes in the combination of temperature and precipitation over the Holocene, which are equivalent to travelling from Minnesota to New England today (as characterized by the climate history at Blood Pond in Fig. 1B). Early in the Holocene, both mean July temperatures and total annual effective precipitation increased substantially (by ∼2-3°C in Fig. 2A and >300 mm in Fig. 2B), but after ca. 7000-5000 YBP, a long-term trade-off between temperature and effective precipitation shaped the regional history. July temperatures declined by ∼1°C, but effective precipitation increased by an additional ∼300 mm (Fig. 2A-B). Similar trends also affected the western portion of *Tsuga*’s range (Calcote, 2003). Because the effective precipitation changes were large, populations tracking their optimal climate conditions would have had to move westward more than northward. Climates once found in central Massachusetts now exist in Minnesota, Wisconsin, Michigan, and western New York (Fig. 1B).

In addition to the long trends, paleoclimate time series also show evidence of 1) abrupt cooling episodes, notably by >1°C at 5400 (5650-5300) YBP in the region of the Labrador Current (GGC30 in Fig. 2A), and 2) multi-century droughts recorded at both inland and coastal areas at 4200–3900, 2900–2100, and 1300–1200 YBP (Fig. 2B). Some changes were spatially heterogeneous, such as at 5800 YBP when water levels fell at the western lake-level site, Davis Pond, but rose at the eastern lake-level site, Deep Pond (Fig. 2B)(Newby et al., 2014; Shuman and Burrell 2017).

The changes were ecologically large. They equal differences among biomes today. Increases in temperature and precipitation after 12,000 YBP compare to moving today from boreal forests in northern Minnesota down through forests with *Tsuga* populations in Wisconsin and Michigan (blue symbols, Fig. 1B). Rapid summer cooling after ca. 5700 YBP was then analogous to shifting eastward to the lowlands of western New York and southern Ontario, or in some cases, coastal southern New England, where *Tsuga* pollen percentages have been low since before European settlement (orange, Fig. 1B). Finally, additional cooling and an increase in effective moisture by 3200 YBP (Fig. 2) equaled moving into the New England highlands where *Tsuga* pollen percentages are high today (black, Fig. 1B).

The climate trajectory of each site thus passed through unique combinations of absolute temperature and precipitation with almost no repetition of earlier combinations (Fig. 5A). Climates of the early Holocene did not overlap with those of more recent millennia (e.g., compare colored symbols representing early-, middle-, and late-portions of the Holocene, Fig. 1B). The differences prevent the early *Tsuga*-climate relationship at any given site from being used to predict later abundance at that site and vice versa. Data from the whole environmental space are required.

**Figure 5.**
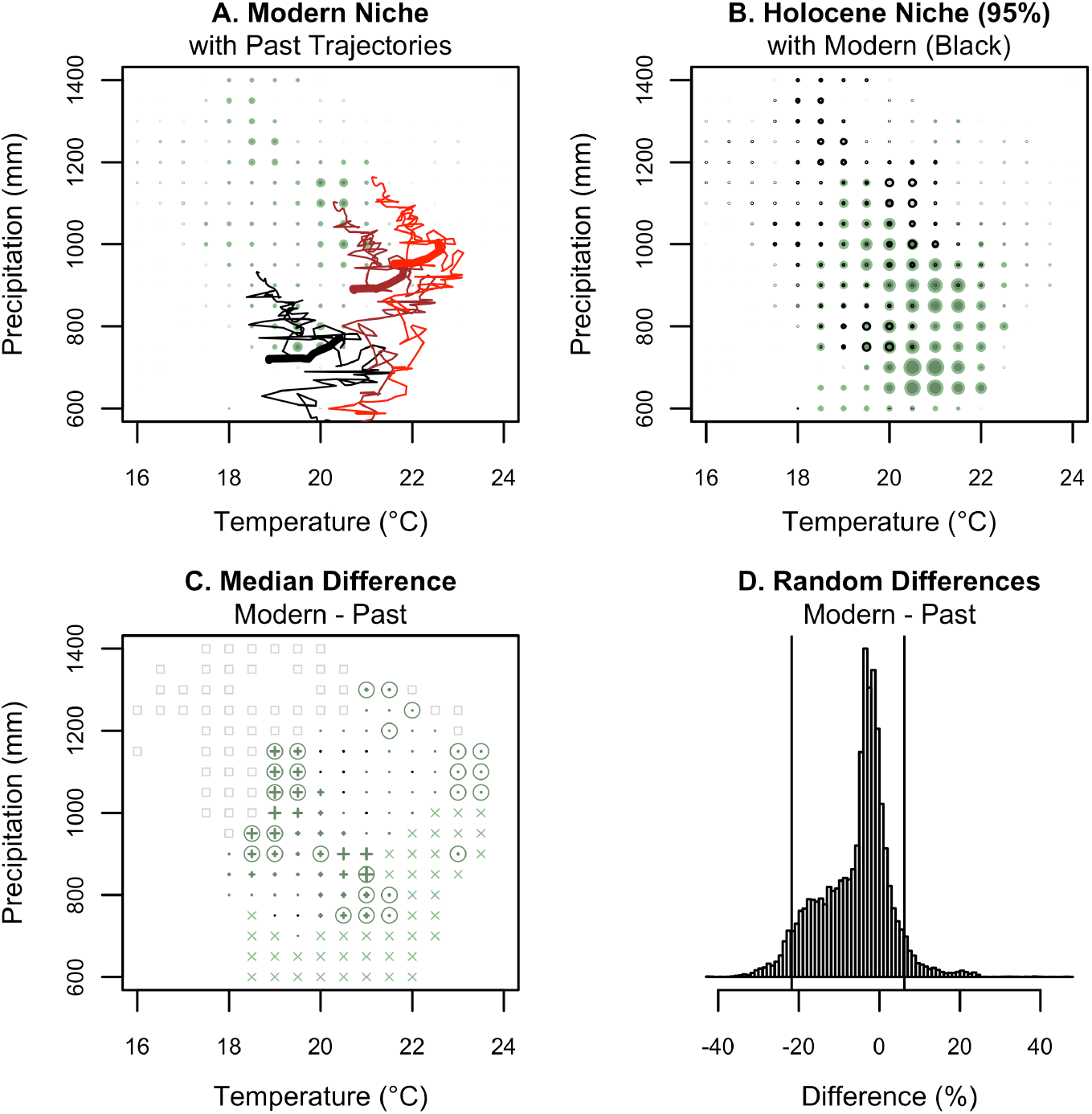
Plots show *Tsuga* pollen percentages today (A), during the Holocene (B), and their differences (C, D) with respect to mean July temperature and annual precipitation and the trajectories of three study sites (colored lines in A). Green circles indicate the upper 95% of the distribution of pollen percentages within 0.5°C and 50 mm windows with the largest symbols representing 50% *Tsuga* pollen. Modern pollen percentages also appear as black circles for comparison with green Holocene values in B. Panel A includes the climate trajectories of Knob Hill (black), Blood (brown), and Spruce ponds (red) with the period of rapid regional cooling at 5600-5300 YBP in bold. The lower end of each thin line represents the oldest samples; the right end of each bold line indicates 5600 YBP and the left end indicates 5300 YBP. Panel C shows the median differences between 1000 random pairs of modern and Holocene samples within each climate window: green plus symbols, scaled to the difference like the circles in A and B, indicate a greater median value during the Holocene than today. The few black symbols indicate the reverse. Circled symbols indicate significant differences. “X” denotes windows with non-zero Holocene medians but a zero median today; gray squares indicate modern climates not experienced by the study sites during the Holocene. The histogram in D shows the differences between all randomly paired samples with vertical black lines depicting the 95% range of differences.

The individual sites followed nearly parallel trajectories through different portions of the modern climate niche (Fig. 5A). Each site began the Holocene where effective annual precipitation was too low for *Tsuga* (like Minnesota today). As temperatures and effective precipitation increased, the sites then moved upward to the right in Fig. 5A and into the area of climate space occupied by *Tsuga* today. Some sites then remained within the climate space represented by the negative (Great Lakes) group of modern *Tsuga* samples (e.g., the trajectory at Knob Hill Pond, black line in Fig. 5A), but others crossed into the space occupied by the positive Appalachian group (e.g., at Blood Pond, brown line in Fig. 5A) or remained at the margins (e.g., Spruce Pond, red line in Fig. 5A).

After ca. 7000 YBP, the locations in climate space shifted up to the left in Fig. 5A because temperatures decreased as effective precipitation increased. The mean slope of the temperature-precipitation correlation after ca. 7000 YBP equaled −97.5 mm/°C, and ensured that *Tsuga*-dominated sites tracked within the ridges of *Tsuga* abundance (e.g., Knob Hill and Blood ponds, black and brown lines in Fig. 5A respectively). When temperatures declined by 5400 YBP (Fig. 2A), however, climate trajectories shifted down to the left in Fig. 5A (bold lines) and cut across rather than along the ridges of sustained *Tsuga* abundance. At this one time in the past 7000 YBP, the climate trajectories for the sites with abundant *Tsuga* moved out of the optimal climate space for high *Tsuga* pollen percentages (Fig. 5A). The addition of a third variable, such as winter temperature, could have shifted the conditions even further from *Tsuga*’s optima, but we lack suitable data to evaluate this possibility.

### 4.2 Niche realizations through time and space

Combining the paleoclimate trajectories inferred from the lake-level and alkenone records with the observed pollen percentages at our eight sites provides a Holocene perspective on the realized climate niche (Fig. 5-6). From this perspective, *Tsuga* pollen percentages also express two parallel maxima in climate space with negative temperature-precipitation correlations (green symbols, Fig. 5B). The reconstructed Holocene niche occupied some warmer and drier climates than the modern niche (compare green versus black symbols in Fig. 5B), and where the two realizations of the niche overlap, fossil percentages were significantly greater than those today in 28 of 85 individual windows in climate space (32.9% of cells in Fig. 5C). Overall, however, the 95% distribution of all modern-minus-fossil sample differences (10.0 to −24.6 percentage points; median: −4.0 percentage points) includes zero and is consistent with many differences deriving from stochastic processes and reconstruction uncertainties (Fig. 5D).

**Figure 6.**
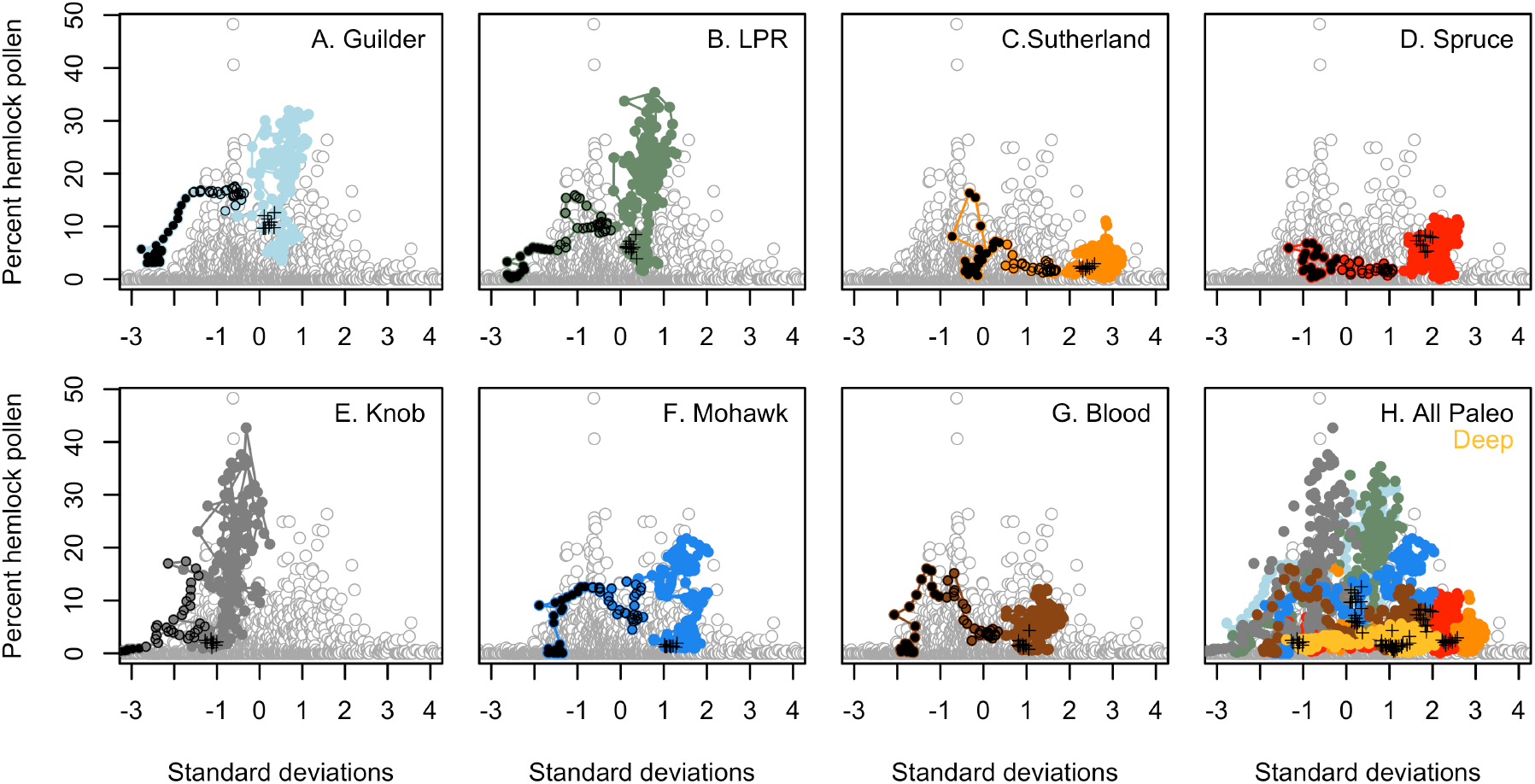
Fossil *Tsuga* pollen percentages from each study site (colored symbols) are plotted with respect to all modern samples (gray) and their climates, represented as standard deviations from the mean climate of all sites where *Tsuga* pollen percentages exceed 1% today (as in Figure 4B). Black circles represent samples older than 9000 YBP; black filled symbols represent >10,500 YBP and black outlined symbols, 10,500-9000 YBP. Black crosses indicate the 500-yr period after the decline from 5200-4700 YBP when percentages were low but retained bimodality (panel H). Because the regional climate became wetter through time (Fig. 2B), the oldest samples plot furthest left (negative departures from the mean) and the youngest samples to the right (positive departures from the mean). Sites shown are Guilder (light blue), Mohawk (dark blue), Sutherland (dark orange), Spruce (red), Knob Hill (dark gray), Little (LPR, dark green), Blood (brown), and Deep ponds (light orange). All sites are shown together in panel H. A supplementary animation shows the realized expression of these relationships by 500-yr time slice.

Over time, the pollen percentages increased or declined in a manner consistent with the bimodality observed in the climate niche today (Fig. 6; Supplementary Animation). The independence of the modern and paleoecological datasets required no alignment of past and modern patterns, particularly early in the Holocene (>9000 YBP, black highlighted symbols in Fig. 6). However, our study sites followed bimodal trajectories within the climate-abundance space delineated by modern data (gray in Fig. 6) and reached two maxima like those observed today: one in association with cool, dry climates (negative departures from mean conditions, Fig. 6) and one with warm, moist climates (positive departures from mean conditions, Fig. 6). All sites experienced minima in abundance when their local climate neared the mean climate of the distribution (zero in Fig. 6).

The trajectories of sites with high percentages tracked within the two modern modes (e.g., Blood and Knob Hill ponds, Fig. 5-6) whereas those with low percentages followed the margins (e.g., Spruce Pond, Fig. 5-6). The amplitudes of change typically declined as sites represented increasingly marginal climates for *Tsuga* (Fig. 6H). As a result, sites like Spruce and Sutherland ponds (Fig. 6C-D) followed bimodal trajectories with low maxima (<10%) about 1-2 sd from the mean combination of temperature and precipitation, whereas sites like Guilder and Little ponds reached high maxima (>15%) at 0.5-1 sd from the mean (Fig. 6A-B). Knob Hill Pond in cool, dry Vermont did not follow a bimodal trajectory, but remained entirely within the climate space occupied by the modern negative (cool, dry Great Lakes) mode (Fig. 6E).

The bimodality in climate space corresponds to peaks in *Tsuga* pollen time series. For example, when *Tsuga* pollen percentages first increased in New England before 9000 YBP, only the negative (cool, dry Great Lakes) mode of the modern distribution was well represented (black circles in Fig. 6; see also the early frames of the Supplementary Animation). These early samples form the earliest peaks in *Tsuga* pollen percentages, like those at Sutherland and Spruce ponds (Fig. 2C)(Maenza-Gmelch, 1997) and elsewhere in the mid-Atlantic region (Zhao et al., 2010). Because the different sites experienced the optimal absolute conditions for peak percentages at different times (e.g., at different latitudes and elevations), the early peaks differ modestly in time (compare black filled versus open circles for >10,500 YBP and 10,500-9000 YBP respectively at sites like Sutherland and Guilder ponds in Fig. 6; see also Fig. 2C).

Low *Tsuga* pollen percentages then followed from ca. 9000-8000 YBP as most sites moved climatically between the modes of the niche (and the youngest black circles plot near zero along the x-axis in Fig. 6). At the same time, however, *Tsuga* pollen percentages first increased at our coldest and driest site, Knob Hill Pond, which did not experience an early peak in abundance (Fig. 2B) and never experienced the climate associated with the modern positive mode (Fig. 6E; black symbols in the Supplementary Animation).

After 8000 YBP, new combinations of temperature and precipitation coincided with renewed increases in *Tsuga* pollen percentages. The changes established the positive (warm, moist) mode in New England where it remains today (positive x-axis values in Fig. 6). When *Tsuga* pollen percentages declined at most sites by ca. 5200 YBP (black crosses in Fig. 6), conditions usually shifted away from each site’s local optimum and back towards the local intra-mode minimum, close to earlier minimum samples from 9000-8000 YBP (compare black circles and crosses in Fig. 6). High percentage samples at Guilder and Little ponds at x = 0 (i.e., above the black crosses in Fig. 6A-B) represent the transient values of the decline itself, which aligns with the intra-mode minimum (Supplementary Animation). After the decline, bimodality remained with peaks in abundance represented by Little, Guilder, and Spruce ponds (crosses in Fig. 6H). As effective precipitation increased after ca. 3700 YBP, the sites re-occupied optimal climates and *Tsuga* pollen percentages correspondingly increased again.

## 5. Statistical models of *Tsuga* pollen percentages

### 5.1 Climate signals

The similarities between modern and past *Tsuga*-climate relationships (Fig. 5-6) could only emerge from persistent and robust associations among the paleoclimate and pollen time series. In fact, the sum of the positive anomalies in the alkenone temperature and lake-based effective precipitation reconstructions (normalized as z-scores) indicate that the sequence of climate changes would have repeatedly favored high *Tsuga* abundance (Fig. 7). The signal includes a sharp rise to peak favorability by 8200 YBP, an abrupt mid-Holocene decline in favorable conditions, a minimum from ca. 5000-4000 YBP, and then a progressive rise in favorability over the last four millennia (e.g., representative examples from Deep and Knob Hill ponds, Fig. 7). Key local details are also similar including short-lived maxima in both the climate signal and *Tsuga* pollen percentages between droughts at Deep Pond at 4350, 3000, and after 1600 YBP (Fig. 7A) and a short-lived minimum associated with cooling at 6250 YBP at Knob Hill (Fig. 7B). Notably, the sharp mid-Holocene declines in the climate-derived time series have different median ages (5150 YBP at Deep Pond and 5550 YBP at Knob Hill), which parallel the observed difference in the median ages of the local *Tsuga* declines: 5150 YBP at Deep Pond and 5500 YBP at Knob Hill. An important area of mismatch relates to differences before ca. 8500 YBP (possibly because this heuristic approach would not account for bimodality in the niche). Based on the coherency of these climate signal models and the fossil pollen data, we proceeded to develop GAMs to quantify the relationships.

**Figure 7.**
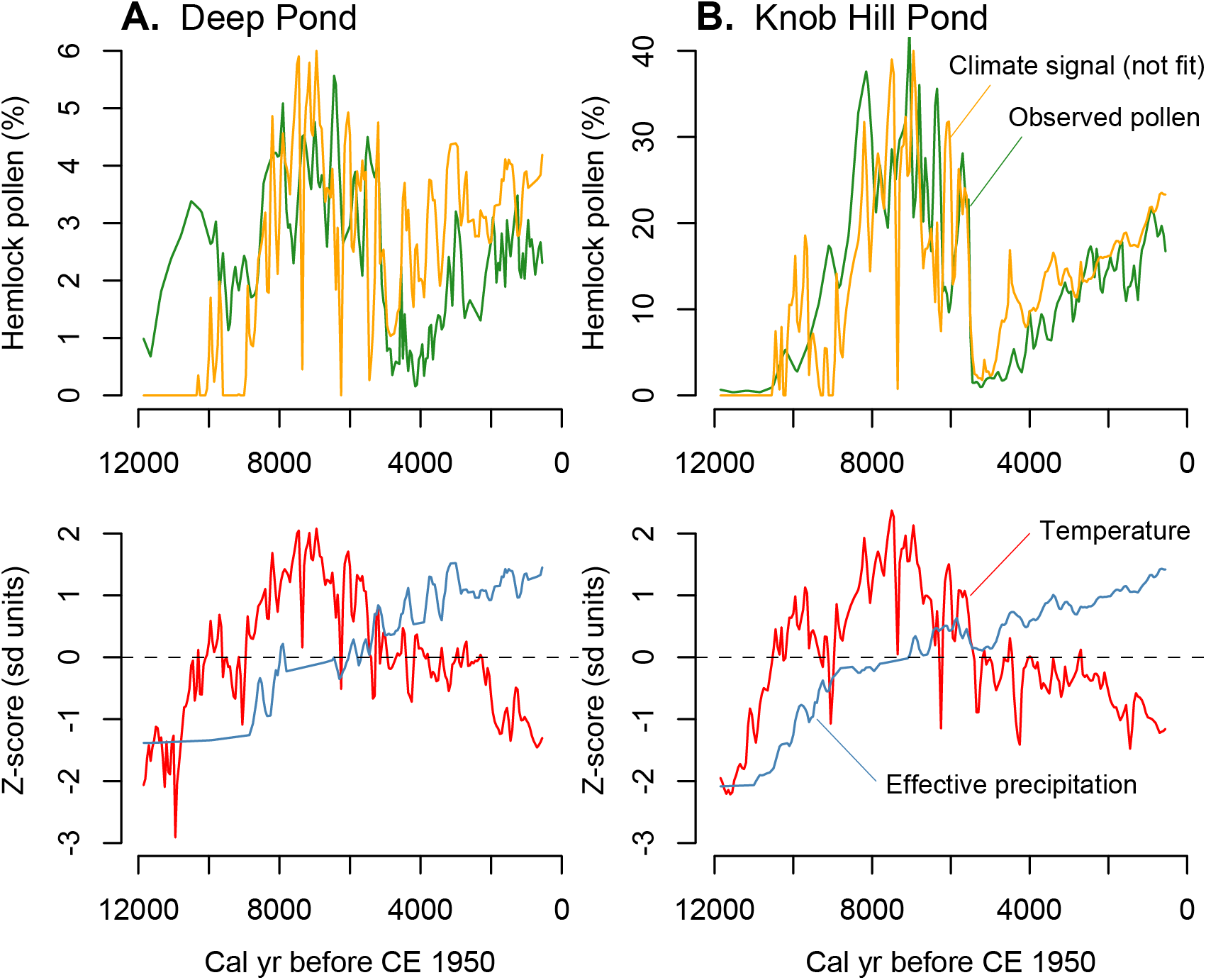
*Tsuga* pollen percentages from A) Deep and B) Knob Hill ponds (green) are shown with a climate index (orange) defined by the sum of the positive departures from the Holocene mean (z-scores) of July temperatures and annual precipitation for each site and scaled to the variance of the pollen data. The positive z-scores represent increases in either temperature or precipitation, like those represented by bold arrows in Fig. 3A, which would favor abundant *Tsuga*. The z-scores of the interpolated temperature (red) and precipitation (blue) series for each site are plotted versus time below.

### 5.2 Generalized additive models of Tsuga abundance

GAMs based on the site-specific estimates of mean July temperatures and annual effective precipitation explain the majority of the variance in the *Tsuga* pollen time series (mean adjusted R^2^ = 0.60; range = 0.29-0.85; Table 2). Root mean squared errors (RMSE) varied from 0.9-5.2 percentage points (Fig. 8-9), and equal 12-15% of the local maxima in *Tsuga* pollen percentages (Table 2). The variance explained ranges from 69% to 85% at sites with high *Tsuga* (>20%) pollen percentages (Guilder, Knob Hill, Little, and Mohawk ponds), and from 29% to 54% at the low abundance sites (Blood, Deep, Spruce, and Sutherland). The variance explained by the GAMs based on the climate reconstructions is significant. It consistently exceeds the 95% range of variance explained by auto-correlated random time series (Fig. 10, Table 2). The null distributions show the potential spurious explanatory power arising from autocorrelated predictor variables and a flexible GAM, but models based on the actual paleoclimate reconstructions have more explanatory power than expected from these factors alone (Fig. 10).

**Table 2.** Model statistics. “Maximum” refers to the maximum percentage of *Tsuga* pollen at each site. “RMSE” is the root mean squared error of each model, and for the regional models refers to the error at the site excluded from the model. The regional model r^2^_region_ represents the variance explained across all other sites, and r_site_ indicates the Pearson correlation coefficient of the observed versus predicted percentages at the excluded site; * indicates p < 0.05 and **, p < 0.0005. The “RMSE/Max” is the proportion of the RMSE to the maximum *Tsuga* pollen percentage at each site. “Converged” represented the percentage of random models that converged without failing.

| Site | Maximum<br>(%) | Random null models |  | Site-specific model |  |  | Regional model with site<br>excluded |  |  |
| --- | --- | --- | --- | --- | --- | --- | --- | --- | --- |
| | | $r^2$<br>(5-95% range) | converged<br>(%) | $r^2$ | RMSE<br>(%) | RMSE/Max | $r^2_{\text{region}}$<br>*Deficient models* | $r_{\text{site}}$ | RMSE<br>(%) |
| Knob Hill | 43 | 0.11-0.57 | 89 | 0.85 | 5.0 | 0.12 | 0.71 | -0.08 | 16 |
| Guilder | 32 | 0.09-0.50 | 90 | 0.69 | 4.5 | 0.14 | 0.70 | 0.14* | 10 |
| Little | 35 | 0.09-0.60 | 92 | 0.76 | 5.2 | 0.15 | 0.73 | 0.23** | 11 |
| *Complete models* |  |  |  |  |  |  |  |  |  |
| Blood | 16 | 0.06-0.37 | 90 | 0.54 | 2.4 | 0.15 | 0.75 | 0.31** | 4.3 |
| Mohawk | 22 | 0.06-0.48 | 85 | 0.70 | 3.2 | 0.15 | 0.75 | 0.65** | 5.6 |
| Deep | 6 | 0.05-0.35 | 90 | 0.43 | 0.9 | 0.14 | 0.70 | 0.60** | 2.1 |
| Sutherland | 16 | 0.03-0.25 | 91 | 0.29 | 1.9 | 0.12 | 0.73 | 0.23** | 3.1 |
| Spruce | 12 | 0.08-0.46 | 90 | 0.54 | 1.8 | 0.15 | 0.73 | 0.13* | 5 |

**Figure 8.**
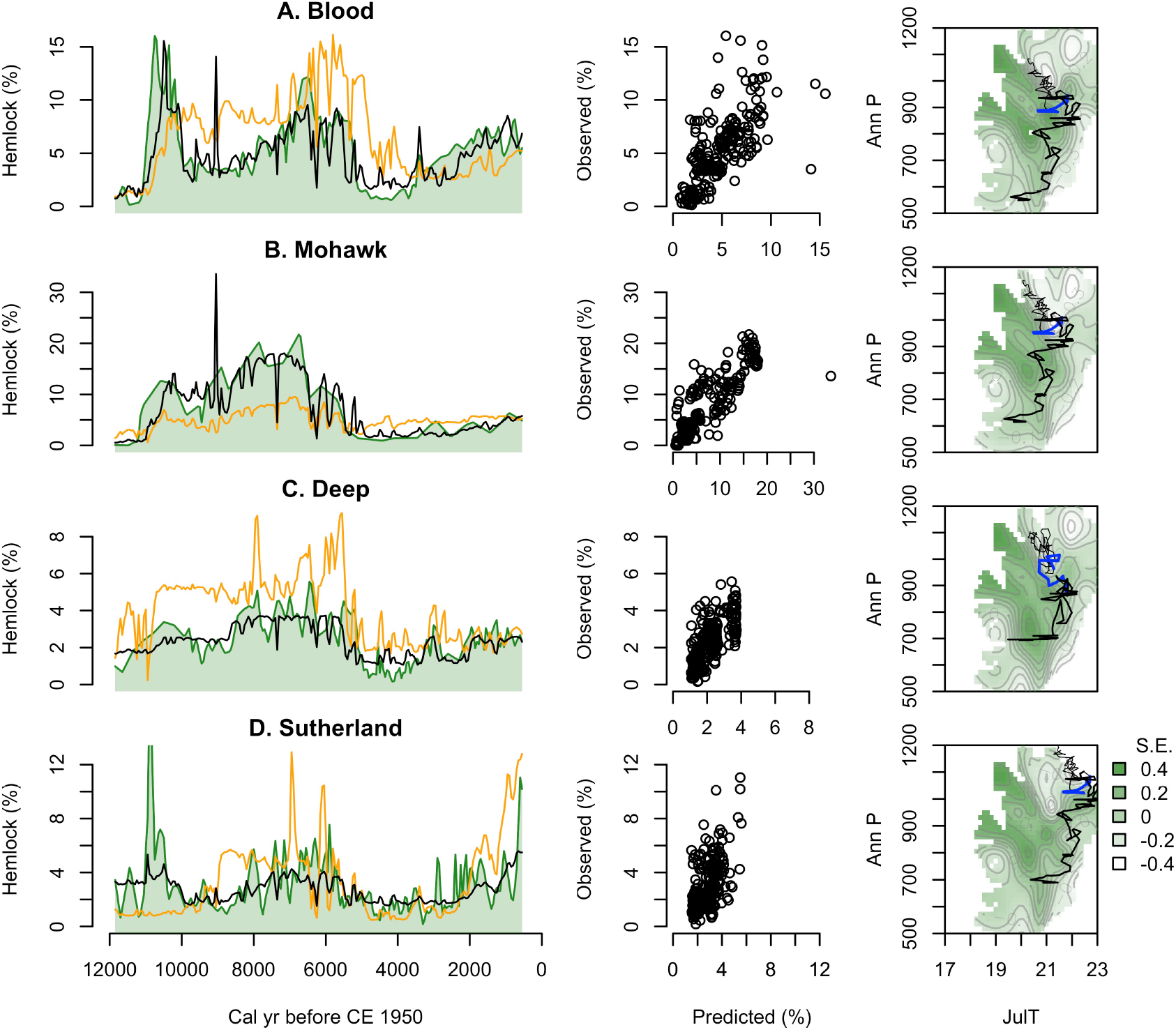
Comparison of observed *Tsuga* pollen percentages with site-specific and regional “leave-one-out” models for Blood, Mohawk, Deep, and Sutherland ponds. Left panels show the observed pollen percentages (in green) plotted versus time with the predicted values from both the site-specific (black) and “leave-one-out” (orange) GAMs. Scatter plots compare the observed and predicted values for each site-specific model. The right-most panels represent the response surface produced by fitting a GAMM to all data from >500 YBP in the region except for the target site. Contours represent increments of 0.1 standard error (S.E.) from the mean; light shades of green represent areas of low pollen percentages and dark shades represent areas of high percentages. Black lines show the climate trajectory of each individual site relative to mean July temperatures (JulT) and annual precipitation (AnnP) with blue line segments representing the period from 5600-5300 YBP associated with the classic *Tsuga* decline. Thin black lines represent the period after 5300 YBP whereas thick black lines represent the period before 5600 YBP.

The regional “leave-one-out” models consistently fit 70-75% of the variance in the data used to develop each model (Table 2). Five models are sufficiently complete that they do not differ meaningfully from each other (right panels, Figs. 8 & 9D) or from the complete dataset (Fig. 5B), but three deficient models do not fully represent the Holocene climate space or the details of the *Tsuga*-climate relationship (right panels, Fig. 9A-C). Overall, correlation between predicted and observed pollen percentages at sites predicted using complete models ranges from 0.23 to 0.65, but those for incomplete models are <0.23 (Table 2). All of the complete models accurately predict a mid-Holocene decline (Fig. 8).

**Figure 9.**
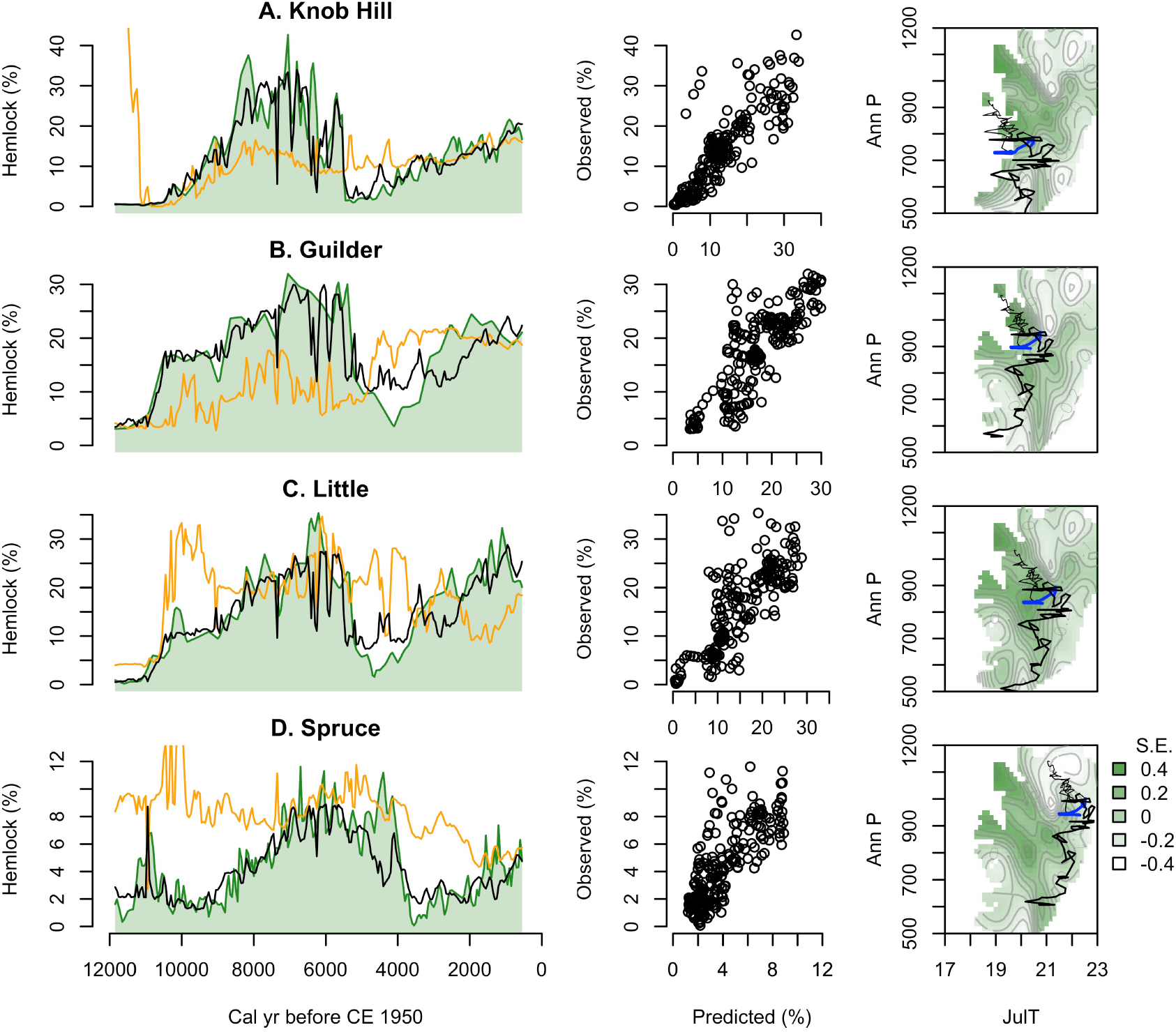
Same as Figure 8 but for study sites with deficient or erroneous regional “leave-one-out” models (Knob Hill, Guilder, Little, and Spruce ponds). Comparison of the surfaces in the right panels with those in Figure 8 or for Spruce Pond in D reveals truncated coverage, gaps, or misrepresentation of the bimodality in the models in A-C that target Knob Hill, Guilder, or Little ponds.

The five complete models individually exclude Blood, Mohawk, Deep, Sutherland, and Spruce ponds. The first three experienced paleoclimates that followed trajectories through well-constrained portions of the model (right panels, Fig. 8A-C), whereas Sutherland and Spruce ponds lie along the margin (Figs. 8D & 9D). In each model, the *Tsuga*-climate relationship is bimodal with the maxima in the pollen percentages expressing negative temperature-moisture correlations as described for modern data (dark green in right panels, Fig. 8). Limited constraints on the random site effects (e.g., soils) cause the predicted and observed mean percentages to differ (orange and green lines in left panels, Fig. 8), but the models accurately predict important features of the pollen percentages at each excluded site including the mid- and late-Holocene maxima and the abrupt mid-Holocene decline. They also accurately represent additional short-lived fluctuations at ca. 4300 and 3000 YBP at Deep Pond (orange lines, left panels, Fig. 8).

Other details are poorly predicted, including the early Holocene maximum at Sutherland Pond (orange line, left panel Fig. 8D). The Spruce Pond model represents the late decline there after ca. 3500 YBP and includes early and mid-Holocene maxima, but the *Tsuga* pollen percentages are commonly over-predicted and an inaccurate second decline is predicted after ca. 2000 YBP (orange line, left panel Fig. 9D). Although the data from Spruce Pond inform the relevant region along the margin of the Sutherland Pond model and vice versa (note the similar trajectories in Figs. 8D & 9D), the models correctly predict large differences in the timing of the decline at the two sites. Otherwise, the predictions for Sutherland are more accurate than for Spruce, especially after 8000 YBP.

The deficient models target three high abundance sites, Knob Hill, Guilder, and Little ponds (Fig. 9A-C). They predict some commonly observed features such as early-Holocene peaks in *Tsuga* pollen percentages, but fail to produce the mid-Holocene decline and do not fully represent the bimodal structure of the *Tsuga*-climate relationship. At Knob Hill Pond, the reconstructed climate at the time of the decline lies outside of the well-constrained model space (blue line in right panel, Fig. 9A) and, consequently, no decline is predicted (orange line in left panel, Fig. 9A). At Guilder Pond, the regional model is similarly incomplete (blue line in right panel, Fig. 9B); the model produces a rapid shift in *Tsuga* abundance in the mid-Holocene, but in the wrong direction (Fig. 9B). The regional model for Little Pond insufficiently represents the minimum between modes in *Tsuga* pollen percentages and contains a plateau of high abundance with no clear bimodality (right panel, Fig. 9C); the oversimplified model prohibits the prediction from declining to a minimum (orange line in right panel, Fig. 9C). In the raw climate-abundance relationships for both Guilder and Little ponds, however, low percentages immediately after the decline align with the intra-mode minimum in the modern climate niche (black crosses, Fig. 6A-B). These GAMs fail, but they should be expected to fail. They are incomplete or inaccurate representations of the climate niche.

## 6. Discussion

### 6.1 A complex climate niche

*Tsuga*’s realized climate niche has two distinctive features, which appear to have existed throughout the Holocene: bimodality and a negative correlation between temperature and precipitation (Fig. 4-6). At any given point in time, our few samples from New England may not fully realize or may misrepresent through under-sampling (alias) structure within the niche (Supplementary Animation), but by sampling across the full range of Holocene climates represented across space and through time, we obtain a more complete view of fundamental or conservative aspects of the niche than otherwise possible. When the full trajectories of the individual sites are evaluated, the bimodality (Fig. 6) and negative temperature-precipitation correlation apparent today (Fig. 5B) remain despite our Holocene analysis excluding samples from <550 YBP. When models represent these major features of the climate niche, they accurately reproduce the Holocene history of *Tsuga* (Fig. 8) despite the many potential sources of analytical error such as climate reconstruction and age uncertainties, extrapolation of climate anomalies from a few sites to a region, and poorly constrained site effects. When models do not resolve the major structures of the niche, they fail to reproduce the history (Fig. 9).

The two persistent elements of the niche must represent interactions among many processes. For example, *Tsuga* germination peaks at 12-17°C with significant limitations below 6°C or above 21°C (Stearns and Olson, 1958; Olson et al., 1959), but drought within the first several years after germination can cause seedling mortality of >80% (Goerlich and Nyland, 2000). At *Tsuga*’s western range limit, sites show a trade off between temperature and precipitation (one high if the other is low) in association with initial *Tsuga* establishment (Calcote, 2003). Thus, both temperature and moisture availability may interact with *Tsuga*’s distribution in multiple ways from the level of seeds to metapopulations to structure the realized climate niche. Other factors, such as biotic interactions (Rooney et al., 2000; Krueger and Peterson, 2006; Witt and Webster, 2010), correlations among the genetic bases of key traits (Etterson and Shaw, 2001), and correlations among important climate variables (Veloz et al., 2012) probably also contribute to this structure.

The regional bimodality with respect to climate does not appear, however, to represent factors like land use or taphonomy (e.g., pollen percentage effects). The minimum in abundance between modes exists across portions of southern Michigan, Indiana, Ohio, western New York, and Ontario, which have not historically supported many *Tsuga* populations; large *Tsuga* populations were not removed by Euro-American forest clearance (Shane, 1991; Clark et al., 1996; Wang et al., 2015; Paciorek et al., 2016; Goring et al., 2016). Likewise, high abundance of other pollen types did not artificially suppress the pollen signal of *Tsuga*, which was inherently low in this region. Instead, bimodality exists because more than one combination of climate conditions appears to favor high abundance. The multiple combinations of dynamics involved, possibly including its ability to simultaneously compete successfully against deciduous tree taxa under different edaphic extremes at landscape scales (Kessell, 1979), could favor high abundance for different reasons in more than one region.

Additionally, different genotypes could play a role in the bimodality if they are adapted to different climates (Davis and Shaw, 2001). A higher frequency of unique alleles in Midwestern than Appalachian populations could hint at different local adaptations (Potter et al., 2012). However, chloroplast DNA differentiation from the main range to outlying populations appears modest (Wang et al., 1997) and the available genetic data indicate similar ancestries of Midwestern and New England populations (Potter et al., 2012).

Local adaptations in *Tsuga* populations are clear, however, and include differences in the optimal temperatures for germination across geographic regions (Stearns and Olson, 1958) and in seedling carbon fixation rates and water-use efficiency (Eickmeier et al., 1975). The later is relevant to the negative temperature-moisture correlation within the niche because moisture stress substantially inhibited carbon fixation in cool-adapted seedlings from northern Wisconsin compared to warm-adapted seedlings from southern Wisconsin. The seedlings associated with warm southern areas retained 52% of their photosynthetic capacity under drought stress compared to only 29% for those from cool northern areas (Eickmeier et al., 1975). Further analyses are required, but these observations indicate that physiology and population genetics may underlie our finding that *Tsuga* only flourished under cool climates if moisture availability was high and warm climates if moisture availability is low (Fig. 4).

The Holocene niche differs in some respects from the one realized today (Fig. 5B-C). Historic land use could well explain why fossil pollen percentages exceeded modern values in 32.9% of the climate space (Fig. 5C)(Fuller et al., 1998; Goring and Williams, 2017). Additional factors, such as a potentially more mild seasonal range in early Holocene New England than in continental Minnesota today, must have also alleviated some current limitations to broaden the niche into warm, dry climates where *Tsuga* does not currently grow (Xs in Fig. 5C). Even with these differences, and considering the various reconstruction uncertainties, the complex Holocene niche depicted by our dataset conforms closely to the one expressed today (Fig. 5-6).

### 6.2 The role of Holocene climate change

The stability of key aspects of the climate niche through time has an important corollary: the reconstructed Holocene climate changes explain the major features of *Tsuga*’s history over the past 12,000 years. If not, the key features of the realized climate niche would not have been stable (Lenoir and Svenning, 2014).

Several lines of evidence conform to the hypothesis that the major changes in *Tsuga*’s abundance derived from the interactions of independent trajectories of temperature and moisture with a multidimensional climate niche (Fig. 3)(Webb III, 1986; Prentice et al., 1991). First, the independent modern and Holocene climate-abundance relationships are similar (Fig. 5-6). Second, heuristic models (Fig. 7) and site-specific GAMs explain a majority of the variance in *Tsuga* pollen percentages (scatter plots & black lines in Fig. 8-9; Table 2), and the *Tsuga*-climate relationships are stronger than null models would have anticipated (Fig. 10). Finally, complete “leave-one-out” GAMs, while imperfect, predict major features of *Tsuga*’s history at the validation sites, including the mid-Holocene decline (orange lines, Fig. 8).

**Figure 10.**
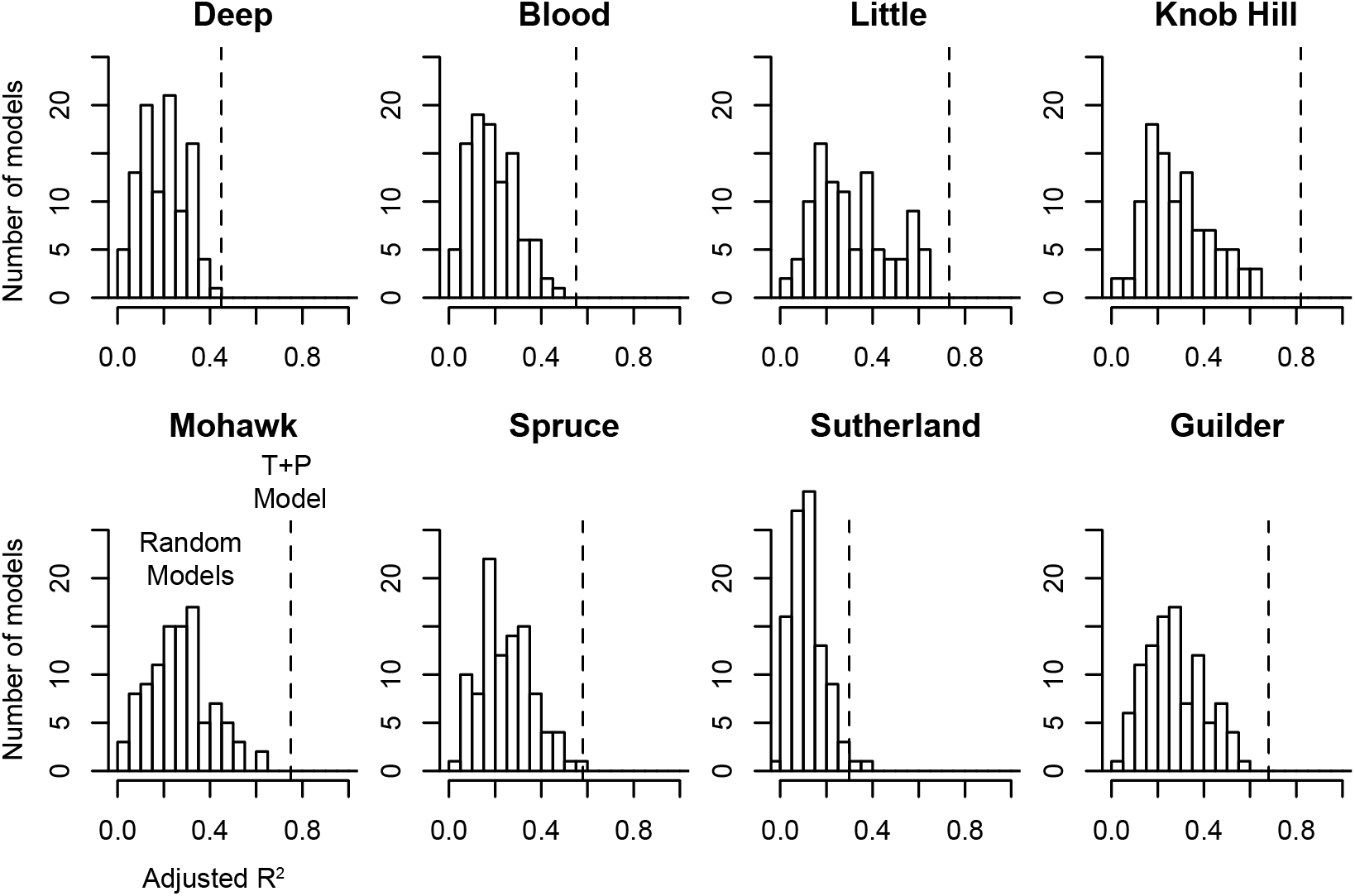
Histograms show the frequency of adjusted (Adj.) R^2^ values for GAMs generated for each site based on 100 pairs of random variables with the same temporal autocorrelation as the reconstructed temperature and effective precipitation time series used in the site-specific GAMs (black lines in Figures 8-9). The adjusted R^2^ for each sites-specific (T+P) model, based on the actual climate reconstructions, is represented by a vertical dashed line.

Davis (1981) ruled out climate as a driver of the *Tsuga* decline, in part, because of a lack of evidence for mid-Holocene climate changes, but such evidence is no longer lacking. The decline emerges as a clear part of the Holocene climate record (Fig. 7), which vegetation history in the region typically tracked (Shuman et al., 2004; 2009; Williams et al., 2002). At finer temporal and spatial scales than considered here, disturbance agents such as forest defoliators may have locally facilitated the mid-Holocene decline (Anderson et al., 1986; Bhiry and Filion, 1996), but the first-order importance of disturbance would represent an unusual, and undetected, exception to persistent regional climate-vegetation relationships (Shuman et al., 2019).

All of the hueristic, site-specific, and complete regional “leave-one-out” models predict abrupt, mid-Holocene declines (Fig. 7-8). Even considering the deficient models (Fig. 9), climate-induced declines should have been common. Furthermore, evidence of insect outbreaks has rarely been found even in locations where they may have been favorably preserved (Oswald et al., 2017) and broadleaf tree taxa susceptible to different diseases and parasites declined synchronously with *Tsuga* in some ecosystems (Foster et al., 2006; Wang et al., 2015). Because the *Tsuga* decline occurred at different times in different places with different absolute climates, the potential role of a single biotic factor that co-varied with climate is also limited (Fig. 8-9). Climate-pollen relationships also did not change during the decline like they did after large disturbances in other conifer-dominated ecosystems (e.g., Calder and Shuman, 2017); model errors here are not systematically correlated with any specific dynamics, times or conditions (Fig. 8-9). Finally, climate-niche interactions also appear to explain similar early Holocene *Tsuga* declines (black circles, Fig. 6; see also Section 6.3).

Overall, the *Tsuga* decline appears no more dependent on disturbance agents than the regional decline of *Picea mariana* (black spruce) and other taxa during rapid climate changes at ca. 11,700 YBP after the cold Younger Dryas period, when biotic disturbance agents are rarely considered (Peteet et al., 1990; Lindbladh et al., 2007; Shuman et al., 2009). The diagnosis retains ambiguities, however. Some of our model validation statistics were low (Table 2) and some residuals large (Fig. 8-9). Other models were inaccurate (Fig. 9). Few sites have been discovered like Spruce Pond where models would predict that conditions favored abundant *Tsuga* during the classic decline period. Despite long-term conservatism of the climate niche (Fig. 5), its observed amplitude collapsed during the decline (black crosses, Fig. 6H). (The models, however, seemingly predict such an outcome by predicting the decline, Fig. 7-8). Solutions to these problems probably involve local edaphic factors, other biotic dynamics, and reconstruction uncertainities as well as the spatial mosaic of climate changes (e.g., drying in the west, but not in the east, Fig. 2B). Other climate variables like winter temperature should also be examined and may be at least as important as any other factor (Calcote, 2003).

Finding that climate history can predict much of *Tsuga*’s past abundance need not, however, represent a climate-versus-ecology dichotomy. The *Tsuga* decline probably represents a mix of multiple proximate dynamics like those during the Younger Dryas that produced abrupt ecological changes up to 100 yrs ahead of the peak rates of climate change (Williams et al., 2002), responses to asynchronous changes in multiple climate variables (Rach et al., 2014), and asynchronous changes across sites (Gonzales and Grimm 2009). Further work should examine how interacting processes from rapid disturbance-induced mortality to long reductions in regeneration played important albeit potentially neutral roles, interchangeably facilitating the climate driven outcomes across large pollen source areas.

### 6.3 Understanding the enigmas of Tsuga’s Holocene history

Our analysis reveals that detailed knowledge of both the multivariate climate history and the complexity of the climate niche can also help to diagnose many of the enigmas of the fossil record of *Tsuga* even if some of the proximate dynamics remain unclear:

#### The complex relationship to Holocene droughts

*Tsuga* abundance depends upon multiple climate variables, and a change in one variable does not always coincide with a change in other (Fig. 2, 7)(Rach et al., 2014). Likewise, the interactions among climate variables can produce more severe outcomes and less intuitive changes than expected from one variable alone (see the example site trajectory as projected with respect to either moisture or temperature alone in Fig. 3A).

Consequently, rapid cooling across the region by 5200±100 YBP overlapped in time with and appears to explain most of the local *Tsuga* declines at 5280±180 YBP (Shuman et al., 2009), even though drought (a decline from the long-term moisture increase spanning the Holocene) did not become widespread until 4925-4575 YBP (Fig. 2C)(Booth et al., 2012; Newby et al., 2014). The cooling interacted with low effective precipitation even before the onset of the multi-century drought because the long-term increase in effective precipitation had not yet reached modern levels (Fig. 2B). The interaction explains why the droughts associated with the *Tsuga* decline were only modest (100 to 150 mm anomalies) compared to the large deviation from modern effective precipitation before ca. 8000 YBP (>300 mm; Fig. 2B). Paleoclimate reconstructions from Michigan and Wisconsin indicate that similar trends extended to *Tsuga*’s western range limit (Calcote, 2003) and thus can explain range-wide changes.

The decline likely had multiple contributing causes (Booth et al., 2012), but they were probably multiple, independently changing climate variables (Calcote, 2003)(Fig. 7). High temperatures were required to sustain high *Tsuga* pollen percentages before the decline because effective precipitation was low and some combination of physiological, developmental, ecological, and population factors underlie a negative correlation between temperature and precipitation in the climate niche (Fig. 4C). Consequently, a sharp drop in temperatures then combined with the persistently lower-than-modern effective precipitation in the mid-Holocene to reduce the potential for high *Tsuga* abundance and cause the decline (Fig. 7). Later, increased effective precipitation enabled *Tsuga* populations to recover because temperatures remained low. Like the bold arrows in Fig. 3A, temperature could cause the decline, but moisture could later move sites back to favorable (cool, moist) portions of the climate niche.

#### Difference in timing of the *Tsuga* decline across sites

Spruce and Sutherland ponds represent two extremes in the local timing of the *Tsuga* decline (3975 and 5600 YBP respectively; Table 1), but differences of centuries also appear to exist between other sites such as Knob Hill and Deep ponds (5500 and 5150 YBP respectively; Fig. 2B). The interaction of three factors explains the differences: 1) spatial variability in temperature change such as rapid early cooling in the north at 5400 YBP that was briefly counteracted by warming at 5250-5200 YBP in the south (Fig. 2A); 2) earlier drought onset in the west than in the east (at 5800 versus 5050 YBP at Davis and Deep ponds respectively; see Newby et al., 2014 for statistical analysis of this asynchrony)(Fig. 2B); and 3) differences in the relationship between *Tsuga*’s climate preferences and the absolute temperatures or effective precipitation at each site (e.g., between high versus low elevation sites such as Sutherland and Spruce ponds).

For example, as the Hudson Highlands cooled and dried after 5650 YBP, *Tsuga* pollen percentages declined at Sutherland Pond, which was higher and wetter than Spruce Pond where *Tsuga* pollen percentages modestly increased until further cooling at 4050 YBP (associated with cooling in the Gulf Stream region recorded by core GGC19, Fig. 2A). The decline at Sutherland Pond (5600 YBP at 380 m elevation) before Spruce Pond (at 3975 YBP at 223 m elevation) provides evidence for a downslope shift in the Hudson Highlands consistent with expectations associated with cooling. The subsequent cooling at ca. 4000 YBP not only facilitated the late *Tsuga* decline at Spruce Pond (Fig. 9H), but also an apparently synchronous *Quercus* decline at Sutherland Pond and a rapid *Betula* increase at Spruce and Sutherland ponds (Maenza-Gmelch, 1997). Likewise, the earlier cooling off Nova Scotia (red line, Fig. 2A) helps to explain why both *Tsuga* and *Quercus* declined on Cape Cod at Deep Pond at 5150 YBP (Foster et al., 2006; Marsicek et al., 2013).

In addition to contributing to variations in the timing of the decline, short-lived climate variations could have also contributed to short-lived declines in *Tsuga* pollen percentages, such as before the classic decline at Knob Hill and other sites (Booth et al., 2012; Oswald and Foster, 2012) and after brief maxima at ca. 4300 and 3000 YBP at Deep Pond (Fig. 8C). Such variability in *Tsuga* pollen percentages may be the result of intrinsic population or ecological dynamics (Williams et al., 2011), but the models show the potential for direct or indirect effects of cold or drought (Fig. 7-8). Because cooling could explain some of the declines, such as after 6250 YBP (Fig. 7-9), drought indicators may not have recorded such events (Booth et al., 2012) and, as with the classic decline, comparisons of the pollen data with only drought indicators could create misleading mismatches.

#### A unique range-wide decline rather than a shift in distribution

At the broad scale, the rapid mid-Holocene decline fits the concept of a “crash” of the realized niche (black crosses, Fig. 6H) because *Tsuga* declined rather than experiencing a range shift (Breshears et al., 2008; Lenoir and Svenning, 2014). The negative correlation between July temperatures and annual precipitation in the climatic niche of *Tsuga* favored declines in all areas of high abundance when climate changes involved positive correlations in the two climate variables (i.e., a reduction in both temperature and precipitation by 4900 YBP; Fig. 5A, bold lines). Because the niche is anisotropic (directionally dependent and not symmetric), climate trajectories perpendicular to the niche orientation limited the number of locations where conditions remained optimal for *Tsuga* populations.

Had the trajectories followed the directionality of the niche (e.g., cooling coinciding with increased moisture as occurred after 2100 YBP), high abundance could have been maintained. When cooling and drying began in the mid-Holocene, however, few fossil pollen sites were warmer and wetter than those in New England, but such environments would have been required to facilitate local increases in *Tsuga* abundance. Spruce Pond, NY, which today is one of the wettest of our study sites and has the highest maximum temperatures (Table 1), may be one of the few sites to meet this criterion. However, other low-elevation sites in eastern New York may also have delayed or missing declines (Ibe, 1982); interpolation of calibrated radiocarbon ages from Heart’s Content Bog, New York, places the decline there at 3700 (3400-4100) YBP, although a hiatus may interrupt the bog stratigraphy (Ibe and Pardi, 1985).

Correlations among temperature and precipitation in the niche were probably as significant as dispersal, biotic interactions, or other factors for limiting the species’ movement because the species could not move into regions with optimal temperatures if precipitation was limiting, and vice versa. The available combinations of temperature and precipitation favored only low abundance (Fig. 7), and interactions with low winter temperatures may have further limited *Tsuga*’s success (Calcote, 2003). Possibly because of antagonistic correlations among the genetic bases for certain plant traits (Etterson and Shaw, 2001), antagonistic relationships within the niche ensured that no large area existed with a suitable combination of temperature and precipitation for abundant *Tsuga* populations from ca. 5200-4000 YBP.

The range-wide collapse has been considered a unique and diagnostic aspect of the *Tsuga* decline (Davis, 1981), but it is similar to several other past biogeographic changes. For example, tree species, such as *Fraxinus* and *Ulmus*, which formed novel (no analog) forest assemblages during the late Pleistocene and early Holocene, also declined across their ranges. Like *Tsuga*, they declined as new combinations of multiple climate variables throughout eastern North America became inconsistent with realizing the highest amplitude portions of their niches (Williams et al., 2001; Veloz et al., 2012). *Tsuga*’s decline was more rapid than the earlier ‘no analog’ declines only because the pace of the relevant climate change was rapid (Fig. 7).

The alignment of *Tsuga*’s niche and subsequent climate trajectories also created a unique range-wide recovery. The recovery parallels the rise of other taxa, such as *Castanea*, which were the climatic opposites of the ‘no analog’ taxa and were initially suppressed by early Holocene conditions. They helped to form new plant communities in the late-Holocene after previously having limited ranges and widespread low abundance (Webb III, 1988). Differences in the species’ climate niches may explain why only *Tsuga* combined the patterns of the no-analog and novel late-Holocene taxa.

Consistent with this interpretation, shifts in the geographic distribution of *Tsuga* at other times during the Holocene coincided with different climate trajectories than those associated with the classic decline (bold versus thin lines in Fig. 5A). For example, after ca. 4000 YBP, a negative correlation in the change in temperature and precipitation followed the directionality of the niche (Fig. 5A). The trend toward cool and wet conditions corresponded with a renewed westward expansion of *Tsuga*’s range beyond Lake Michigan (Davis et al., 1986). *Tsuga* colonized new western sites that were dry where it was warm and wet where it was cool (Calcote, 2003), and thus, tracked a negative correlation between temperature and precipitation like that observed in the climate niche (Fig. 5). Midwestern sites had been both warmer and drier than New England due to their continental position. For this reason, they would have occupied the lower right of Fig. 5 for much of the early Holocene, and would not have been initially suitable for *Tsuga* until conditions cooled and became wetter than before, especially after ca. 3000 YBP (up and left in Fig. 5A).

#### Early-Holocene peaks and declines in only some records

The interaction of the climate trajectories with the complexities of the niche also explains brief peaks in *Tsuga* pollen percentages before ca. 9800 YBP (Zhao et al., 2010; e.g., at Sutherland and Spruce ponds, Fig. 2B). When *Tsuga* first increased in New England, the climate trajectories did not align with the anisotropy of the niche (Fig. 5A). Based on the available paleoclimate data, a rapid increase in effective precipitation corresponded with warming, and thus, cut across the alignment of *Tsuga*’s maximum abundance. Consequently, many first increases were short lived. The increases also correspond to the negative mode of the distribution, dominant in the Great Lakes region today (black circles before 9000 YBP in Fig. 6). Some sites like Little and Guilder ponds (Fig. 6A,F) experienced these conditions at ca. 10,500-10,000 YBP whereas other sites like Sutherland Pond (Fig. 6C) reached the same absolute conditions and peak percentages earlier (Fig. 6). Percentages declined after ca. 10,000 YBP as the climate trajectories crossed out of the climatic region of the negative mode and into the trough in abundance between modes (see the transition from black outlined to colored symbols in Fig. 6, which marks 9000 YBP). Percentages then increased again as the trajectories moved into the climate space of the positive mode dominant in New England today (Fig. 5-6, Supplementary Animation).

The absence of an early peak at Knob Hill Pond (Fig. 7-9) supports this interpretation. Combinations of temperature and effective precipitation at the site persistently remained like those associated with the negative (Midwestern) group of populations today (black, Fig. 6E), and *Tsuga* abundance at Knob Hill increased only after 9800 YBP as it declined from the early peaks in southern New England (Fig. 8A, see also black symbols in the Supplementary Animation). The increase in Vermont, therefore, could represent part of the northwestward shift in populations that ultimately spread the negative group into the Midwest. As a result, the anisotropy of the niche helps to explain the mid-Holocene decline, but the bimodality in the niche explains the early peaks and declines of *Tsuga* before ca. 9800 YBP (Zhao et al., 2010).

### 6.4 Considering multivariate climate change

The relationships in our data indicate that understanding vegetation biogeography at the time scale of the Holocene requires a nuanced and quantitative view of climate history. Multiple aspects of climate changed over this time period. We have not fully constrained that history here (e.g., winter temperatures, seasonal precipitation rates, and the frequencies of extreme events are among the unconstrained variables), but reconstructions of two key variables (summer temperature and effective precipitation) explain much of the variation in the pollen data without evoking processes that were disconnected from climate. Consequently, multivariate climate change may be a primary ecological force at spatiotemporal scales larger than centuries and landscapes. Alternative hypotheses cannot ignore the dynamic climate history, and interpretations must consider that incomplete reconstructions of the paleoclimates (e.g., missing variables, non-linear indices, records of aquatic rather than terrestrial conditions) can emphasize misleading relationships between climate and vegetation. Our results confirm that some paleoecological dynamics, interpreted as a function of non-climatic factors (e.g., diseases, dispersal lags), may have involved climate in important ways that become evident once more than one climate variable is considered.

## 7. Conclusions

Holocene biogeography depends as much on climate as on ecology. The Holocene history of *Tsuga canadensis* reveals that interactions of multivariate climate changes with species’ climate niches probably acted as a first-order control on vegetation history. Independent Holocene and modern realizations of *Tsuga*’s niche share important features and indicate that the relationships between climate and *Tsuga* abundance remained stable through time. Consequently, models of the niche based on independent reconstructions of temperature and precipitation explain 29-85% of the variance in *Tsuga* pollen percentages including the rapid, range-wide decline of *Tsuga* at ca. 5700-5000 YBP. When regional “leave-one-out” models failed to reproduce this history, they did so because they did not completely or accurately represent the major features of the climate niche, such as its biomodality.

The *Tsuga* decline has been attributed to non-climatic factors (Davis, 1981; Booth et al., 2012), but our reconstructions indicate that it coincided with rapid regional cooling when effective precipitation was lower than today. Because *Tsuga*’s climate niche contains a negative correlation between optimal July temperatures and annual precipitation, cooling would have had to coincide with increased precipitation to sustain high abundance. Instead, drought associated with the rapid cooling prevented optimal conditions from developing at most sites until effective precipitation increased in later millennia. While undetectable biotic factors almost certainly played a role in *Tsuga*’s history, we found no direct evidence of such dynamics at the spatial and temporal scales represented by wind-dispersed pollen. Sustained bimodality in the niche further clarifies that many pollen records include an early peak in *Tsuga* abundance before ca. 9800 YBP because the region briefly experienced climate conditions consistent with the Great Lakes mode of *Tsuga* abundance today. As a result, recent forest declines driven by climate changes have precedents in both early- and mid-Holocene declines of *Tsuga*, but recent declines of other taxa (e.g., *Castanea, Ulmus*) attributable to exotic diseases, insects, and other factors do not.

## Supporting information

Supplemental animation

## Acknowledgements

Funding for this project was provided to BS, WWO, and DRF from the National Science Foundation Ecosystem Science Program (DEB-1146297, DEB-1146207). We also benefited from thoughtful comments on the manuscript from T. Webb, J. Williams, M. Fitzpatrick, E. Currano and three anonymous reviewers. Data were obtained from the Neotoma Paleoecology Database (http://www.neotomadb.org), and the work of the data contributors and the Neotoma community is gratefully acknowledged.

## Figure captions

### Supplementary Animation

Plots of *Tsuga* pollen percentages along the axis of temperature-precipitation interaction as in Figure 6 shown by 500-yr time slice. Gray symbols in all panels represent modern values, but colored symbols indicate fossil samples from each site during each 500 yr period. Sites shown are Guilder (light blue), Knob Hill (black), Little (dark green), Mohawk (dark blue), Blood (brown), Deep (light green with black outline), Sutherland (orange), and Spruce ponds (red).

## Appendix 1: Data and Method Details

### Age models

We use the published age-depth relationships in calendar years for all study sites and records, and have made no chronological modifications. In this way, the data are presented as one possible distribution in time allowed by the radiocarbon age uncertainties associated at each site. When we discuss apparent agreement in timing of changes across records, we do so when the mean ages are similar and when even a formal analysis (Parnell et al., 2008) would not be able to rule out synchrony; fine-scale asynchrony within the age uncertainties (years to decades) can never be ruled out with the existing data. When we discuss differences in ages, we have used a formal analysis of synchrony based on the differences between 1000 random samples from the age uncertainty distribution, estimated using *bchron* (Haslett and Parnell, 2008), for each event to test whether the difference was significantly different from zero (Parnell et al., 2008). To enable comparisons across sites, all data were first linearly interpolated to even 50-year time steps. Ages of abrupt changes are based on the peak rates of change measured over 100-yr steps in the interpolated data. We focus on the interval from 550-11850 YBP because of the temporal limitations of the available temperature data (Sachs, 2007).

### Paleoclimate records

The temperature time series used in our analyses represent growing-season SSTs off the coasts of Nova Scotia (core OCE326-GGC30; Sachs, 2007) and Virginia (core CH0798-GGC19; Sachs, 2007), which we assume to also represent regional trends in mean July temperatures of this coastal region. Data from Nova Scotia derive from the region of the Labrador Current, which supplies the Gulf of Maine to the east of New England; data from Virginia derive from the region of the Gulf Stream, which transports heat north along the southern coast of New England. The temperature anomalies are interpolated to the locations of the pollen records based on site latitude to account for mixing of the northern and southern temperature signals; the results build on the high degree of spatial autocorrelation expected for temperature across the region.

Following Shuman and Marsicek (2016), each SST reconstruction was linearly detrended to account for a large trend in the data (e.g., equal to cooling of >5°C since 11,000 YBP in core GGC30), which may be driven by non-climatic controls on alkenone production (Prahl et al., 2006). Without detrending, New England temperatures during the cold Younger Dryas interval (before 11,700 YBP) and other parts of the Pleistocene would have as high as those in modern day North Carolina, which is inconsistent with terrestrial stable isotope records (Huang et al., 2002; Kirby et al., 2002; Hou et al., 2006; Zhao et al., 2010) and the region’s ecological history (e.g., boreal forests in Massachusetts at 12,000 YBP). The detrended signal (Fig. 2A) aligns well with elevational changes in tree species distributions known from plant macrofossils in the White Mountains, New Hampshire and the Adirondacks, New York (Spear, 1989; Jackson and Whitehead, 1991; Spear et al., 1994; Shuman et al., 2004) and with pollen-inferred temperatures for the region (Webb III et al., 1993; Marsicek et al., 2013). The detrended SST records also contain multi-century variability that correlates significantly with variations in the independently derived effective moisture records (Shuman and Marsicek, 2016).

Effective precipitation (precipitation minus evapotranspiration) was derived from reconstructed lake-level changes at Davis Pond in western Massachusetts (Newby et al., 2011, 2014) and Deep Pond on the southern coast of Cape Cod, Massachusetts (Marsicek et al., 2013; Newby et al., 2014). At each site, transects of sediment cores were used to constrain past shoreline position, which we converted to effective precipitation using a simple water budget model for each lake and watershed (Marsicek et al., 2013; Pribyl and Shuman, 2014). Interpolation of the effective precipitation signals from Deep and Davis ponds to the location of the pollen records was carried out based on longitude because of the east-west orientation of the available records (Fig. 1A). The timing of changes in each record is constrained by >3 cores and 31-53 calibrated radiocarbon ages (Newby et al., 2014). As with the pollen and alkenone-derived SST data, no modifications were made to the published age-depth relationships, which were derived using *bchron* (Haslett and Parnell, 2008; Newby et al., 2014).

### Generalized Additive Models (GAMs)

To evaluate the variance in the pollen data explained by the climate reconstructions, we use generalized additive mixed models. They were applied after the pollen percentages were square-root transformed to ensure a normal distribution. The GAMs, thus, take the form *Tsuga*^*0.25*^ = function (July Temperature, Annual Precipitation), and were applied using a tensor product smooth using a cubic regression spline, Gaussian error distribution, and an identity link function in the *mgcv* package in R (Wood, 2006b; Wood, 2011; R Core Development Team, 2017). We used analysis of variance (ANOVA) to compare various models such as those with additive versus interactive predictors, spline versus tensor product smooths, cubic regression versus thin plate regression for each marginal basis, and the inclusion of site as a random effect; the model choices represented the lowest residual degrees of freedom.

To further ensure that the GAMs do not represent spurious correlations common to autocorrelated time series (Granger and Newbold, 1974), we also generated 100 random time series with the same temporal autocorrelation structure as observed in temperature and moisture reconstructions. The approach was inspired by analyses of significance in paleoenvironmental reconstruction by Telford and Birks (2011), and random number series were generated using the function, *gstat*, in R.

### Sources of error

Uncertainty in our analyses stems from factors such as the limits to how well the existing paleoclimate data represent the region, the age uncertainties of the records, the simplified linear interpolation of the climate reconstructions, and the detrending of the SST reconstructions. The alternative GAMs generated using random time series (Fig. 10) indicate that the climate reconstructions must have captured accurate, non-random climate signals to suitably predict the pollen percentages through time even if some errors in absolute climates or ages exist. The long-term negative correlation between temperature and precipitation, as well as the abrupt changes in the SST record at ca. 5500 YBP, exist in the data whether or not we detrend the SSTs. The major differences between the detrended and non-detrended temperature schemes center on the period before ca. 7000 YBP. Therefore, the role of temperature in the *Tsuga* decline is unlikely to derive from our treatment of the temperature data.

The roles of winter temperatures or seasonal precipitation have not been well constrained and could also modify the specific climate effects detected here. Covariance of climate variables could have translated winter signals to either summer temperature or annual precipitation in our models. Likewise, ocean temperatures, likewise, may not represent the regional air temperatures (although see Shuman and Marsicek 2016). Errors could also arise from our assignment of absolute temperature and precipitation values based on different base periods in the 20^th^ century, but sensitivity analyses indicate that the major patterns discussed here are not substantially affected by using a different base period.

